# Evolutionary origin of sex differentiation system in insects

**DOI:** 10.1101/2021.08.02.454784

**Authors:** Yasuhiko Chikami, Miki Okuno, Atsushi Toyoda, Takehiko Itoh, Teruyuki Niimi

**Author notes:** Corresponding author: Teruyuki Niimi.

## Abstract

The evolution of the functionality of genes and genetic systems is a major source of animal diversity. Its best example is insect sex differentiation systems: promoting male and female differentiation (dual-functionality) or only male differentiation (single-functionality). However, the evolutionary origin of such functional diversity is largely unknown. Here, we investigate the ancestral functions of *doublesex*, a key factor of insect sex differentiation system, using the apterygote insect, *Thermobia domestica*, and reveal that its *doublesex* is essential for only males at the phenotypic level, but contributes to promoting female-specific *vitellogenin* expression in females. This functional discordance between the phenotypic and transcription-regulatory levels in *T. domestica* shows a new type of functionality of animal sex differentiation systems. Then, we examine how the sex differentiation system transited from the single-functionality to the dual-functionality in phenotypes and uncover that a conserved female-specific motif of *doublesex* is detected in taxa with the dual-functional *doublesex*. It is estimated that the role of the sex differentiation system for female phenotypes may have evolved through accumulating mutations in the protein motif structures that led to the enhancement of its transcription-regulatory function.

## Introduction

Sex is a fundamental principle in animal reproduction and is shared among almost all animals. The differences between males and females are a large source of diversity in mating systems, species, traits, and ecological dynamics in Metazoa (*Darwin, 1871*; *Fryzell et al., 2019*; *Geddes and Thomson, 1889*). The universality of sex suggests that sex is an ancient feature of metazoans. On the other hand, recent studies in various animals have revealed that regulatory systems for producing sex change rapidly during animal evolution (*Bachtrog et al., 2014*; *Beukeboom and Perrin, 2014*; *Herpin and Schartl, 2015*). For example, genes in the sex determination/sex differentiation systems are more likely to change upstream in the gene cascade than downstream (*Bopp et al., 2014*; *Wilkins, 1995*). In eutherian mammals such as mice and humans, a transcription factor called *Sex-determining region Y* (*Sry*) acts as a ‘master regulator’ of sex determination pathway, in which *Sry* induces expression of a transcriptional factor *doublesex and mab-3 related transcriptional factor 1* (*dmrt1*) in males through chain reactions of transcription factors during embryogenesis (*Gubbay et al., 1990*; *Koopman et al., 1991*; *Miyawaki et al., 2020*; *Sinclair et al., 1991*). In contrast, in medaka fish, *Oryzias latipes*, a paralog of *dmrt1, dmy* (*DM-domain gene on the Y chromosome*) instead of *Sry* promotes male differentiation via a gene cascade inducing *dmrt1* (*Matsuda et al., 2002*; *Nanda et al., 2002*). The diversity in gene repertoires composing sex differentiation systems has also been found in arthropods (e.g., *Sharma et al., 2017*; *Suzuki et al., 2008*; *Xu et al., 2017*).

Diversity of sex differentiation systems has referred to differences in gene repertoires among species or populations. At present, many evolutionary scenarios have been proposed to explain the evolutionary transition of gene repertoires in sex differentiation systems: e.g., neo-functionalization or sub-functionalization via gene duplication (e.g., *Chandler et al., 2017*; *Hattori et al., 2012*; *Hasselmann et al., 2008*; *Matsuda et al., 2002*; *Sharma et al., 2017*; *Yoshimoto et al., 2008*), positional exchange within the cascade via feedback loops (e.g., *Myosho et al. 2012*; *Myosho et al, 2015*; *Smith et al., 2009; Takehana et al., 2014*), and functional shifts via accumulation of mutations in *cis*- or *trans*-elements (e.g., *Kamiya et al., 2012*; *Sato et al., 2010*) (reviewed in *Beukeboom and Perrin, 2014*). Currently, most of the diversity of sex differentiation systems is explained by these well-understood scenarios for changes in gene repertoires. However, recently, a difference in sex differentiation systems without swapping gene repertoires has been discovered from pterygote insects.

Pterygote insects exhibit tremendous sexual dimorphisms in the head, abdomen, wings, and so on, realizing a complex mating strategy. Most of the sexual differences are produced by a sex differentiation system that uses a transcription factor *doublesex* (*dsx*, a homolog of *dmrt1*) as a bottom factor (*Kopp, 2012*; *Verhulst and van de Zande, 2015*). Studies on Aparaglossata (holometabolan insects excluding Hymenoptera) have shown that *dsx* promotes both male and female differentiation via the sex-specific splicing (*Burtis and Baker1989*; *Gotoh et al., 2016*; *Hildreth, 1964*; *Ito et al., 2013*; *Kijimoto et al., 2012*; *Ohbayashi et al., 2001*; *Shukla and Palli, 2012*; *Xu et al., 2017*). Recent studies showed that *dsx* promotes only male differentiation in several hemimetabolan and hymenopteran insects, even though *dsx* has sex-specific isoforms (*Mine et al., 2017*; *Takahashi et al., 2021*; *Wexler et al., 2015*; *Zhuo et al., 2018*). Thus, there is a difference in outputs in the sex differentiation systems in insects: promoting only male differentiation (single-functionality) or both male and female differentiation (dual-functionality). It is suggested that the output of the insect sex differentiation systems transited from the single-functionality to the dual-functionality (*Wexler et al., 2019*). However, it remains largely unclear how the difference in the output evolved independently of changes in gene repertoires and what drove such a transition (*Hopkins and Kopp, 2021*). Also, it is unidentified the ancestral roles of *dsx* isoforms in insects.

The evolutionary origin of the sex differentiation system in insects is ambiguous by the inability to estimate the status of the common ancestor of Insecta. All of the previous studies examining the *dsx* functionality have been limited to pterygote insects or crustaceans and chelicerates. In chelicerates and crustaceans, *dsx* has no sex-specific isoforms (*Kato et al., 2011; Li et al., 2018; Panara et al., 2019; Pomerantz et al., 2015*). We are currently forced to compare the status of crustaceans with that of pterygote insects, resulting in a large gap in phylogenetic mapping. Furthermore, previous reports of the single-functionality have been based mainly on the sexual differences acquired or complicated by each taxon in hemimetabolan insects. Therefore, it remains possible that the single-functionality reported so far results in a secondary loss of roles in female differentiation.

To address these issues, it is necessary to examine the function of *dsx* in taxa that retain the sexual traits of the common ancestor of Insecta and that emerged between the crustaceans and Pterygota. To this end, we focused on the firebrat *Thermobia domestica* (Zygentoma) belonging to Zygentoma, the sister group of Pterygota (*Misof et al., 2014*). The sexual traits of *T. domestica* are restricted to the female simple ovipositor and the male ‘penis’ that is not aedeagus. These sexual traits mirror the ancestral state of insects (*Beutel et al., 2017*; *Boudinot, 2018*; *Emeljanov, 2014*; *Kristensen, 1975*; *Matsuda, 1976*). Here, to examine the exon-intron structure of *dsx*, we decoded the genome of *T. domestica*. Then, we developed the technology to effectively inhibit gene function during postembryonic development and investigated the roles of *dsx* for sexual traits, gametogenesis, and transcriptional regulation, and compared them with other insects.

## Results

### Molecular evolution of Doublesex in Pancrustacea

First, we examined the relationship among *dsx* homologs in animals (Figure 1–figure supplement 1, Table 1) and revealed that the pancrustacean transcription factor *doublesex* (*dsx*) occurs in four clades: crustacean Dsx, Entognatha Dsx, Insect Dsx clade 1, and clade 2. (Figure 1A, B). *Thermobia domestica* has two *dsx* genes, belonging to different clades (Insect Dsx clade 1 and 2) (Figure 1–figure supplement 2). We considered the gene belonging to Insect Dsx clade 1 to be an ortholog of the pterygote *dsx*, since this clade contains Dsx of pterygote insects including *Drosophila melanogaster*. We thus named the gene in Insect Dsx clade 2 *dsx-like* to distinguish it from *dsx*. Insect Dsx clade 2 contained *dsx-like* in Zygentoma as well as Ephemeroptera and Phasmatodea (Figure 1B), indicating that *dsx* was duplicated before the emergence of the Dicondylia (= Zygentoma + Pterygota) and was lost repeatedly in pterygote taxa. This fact supports that *T. domestica* retains the ancestral state for *dsx* related genes’ contents. Since these results suggested the involvement of the gene duplication in the functional evolution of *dsx*, we also analyzed the expression and function of *dsx-like*.

**Figure 1.**
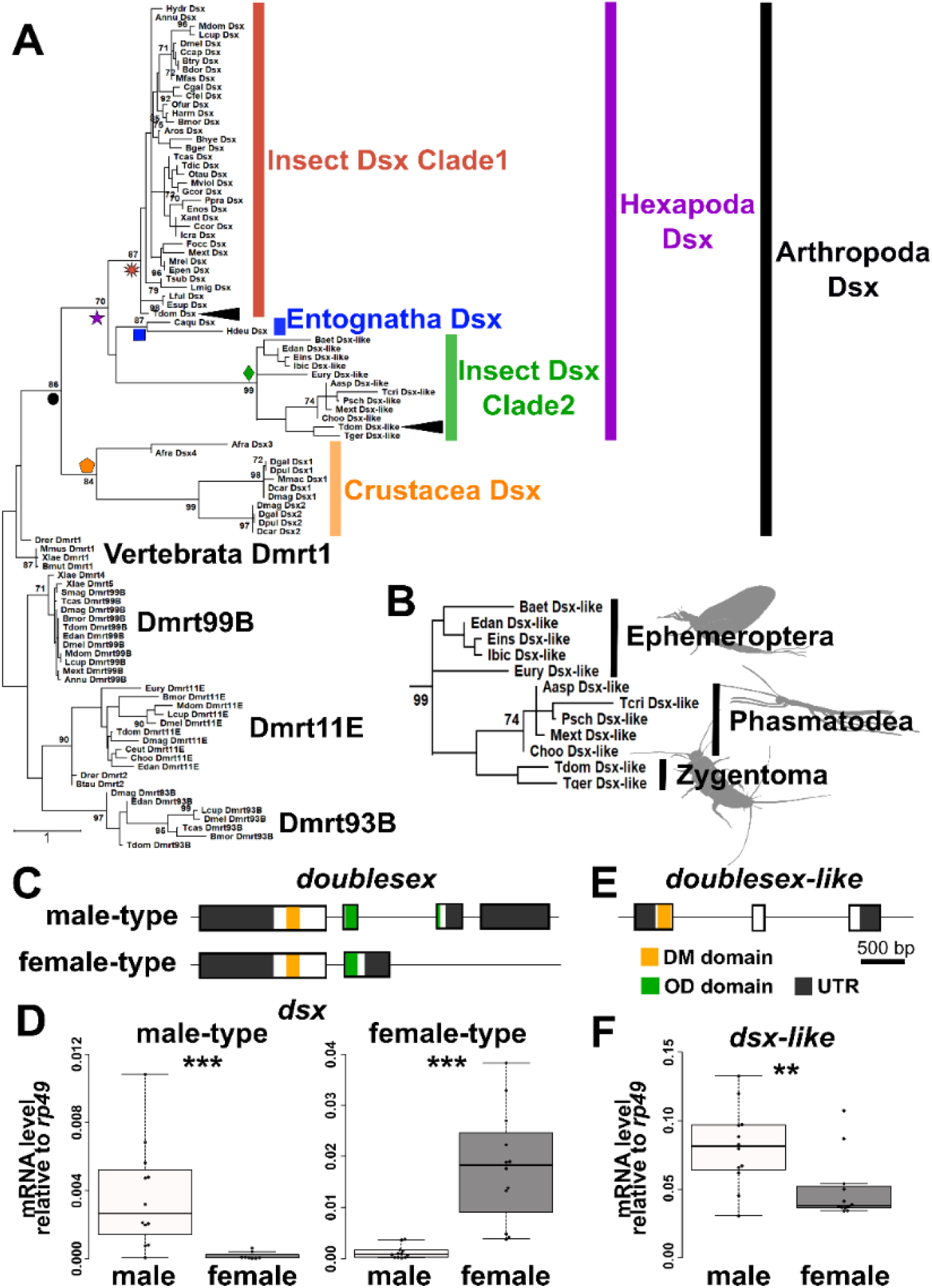
Molecular evolution and features of Doublesex in *Thermobia domestica*. (A to B) Molecular phylogeny of Doublesex and Mab-3 related transcriptional factors (DMRT) (A) and enlarged view of insect Dsx Clade2 (*dsx-like* clade) (B). The numerical value on each node is the bootstrap supporting value. Bootstrap values < 70 are not shown. The node of each clade is indicated by colored shapes: black circle, Arthropoda Dsx; orange pentagon, Crustacea Dsx; purple star, Hexapoda Dsx; red sunburst, Insect Dsx Clade1; green diamond, Insect Dsx Clade2; blue square, Entognatha Dsx. (C to F) Molecular features of *dsx* (C and D) and *dsx-like* (E and F). (C and E) indicate exon-intron structures of *dsx* and *dsx-like*, respectively. (D and F) show mRNA expression levels of *dsx* and *dsx-like*, respectively. Each plot is a value of an individual. Sample size is listed in Table 2. Results of Brunner–Munzel tests are indicated by asterisks: \*\**P*<0.01; \*\*\**P*<0.001 and are described in Table 2.

**Table 1.**
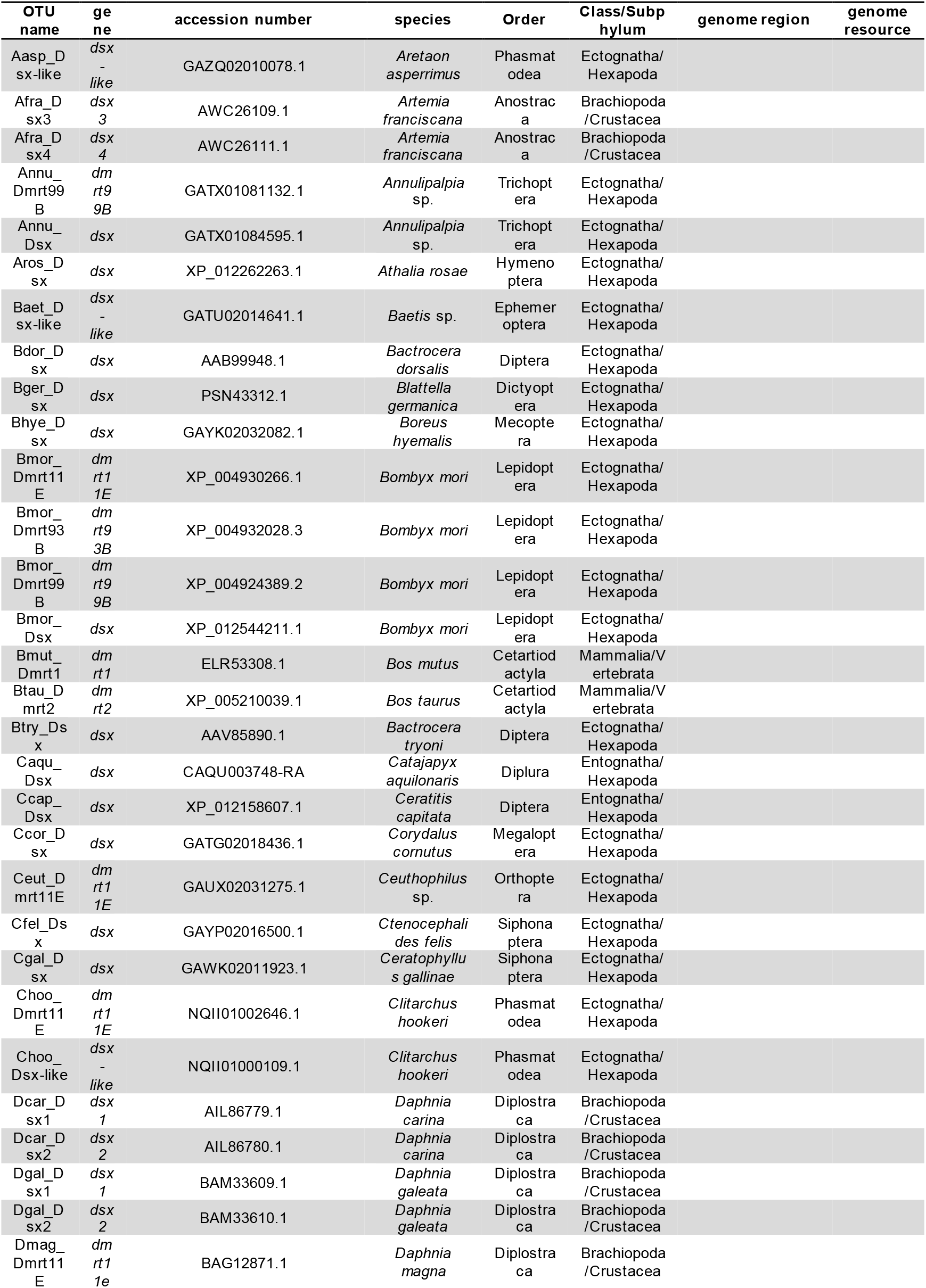
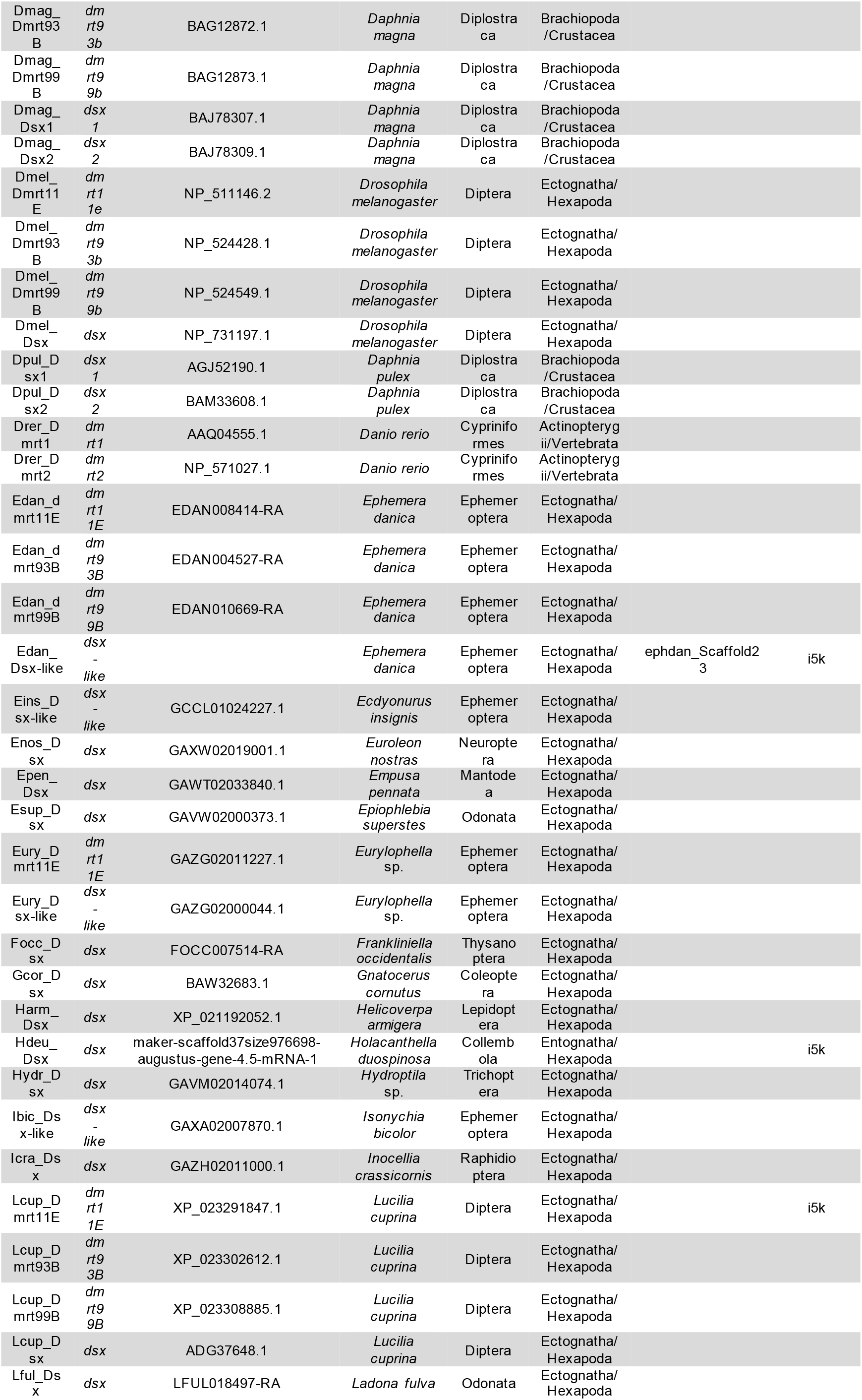
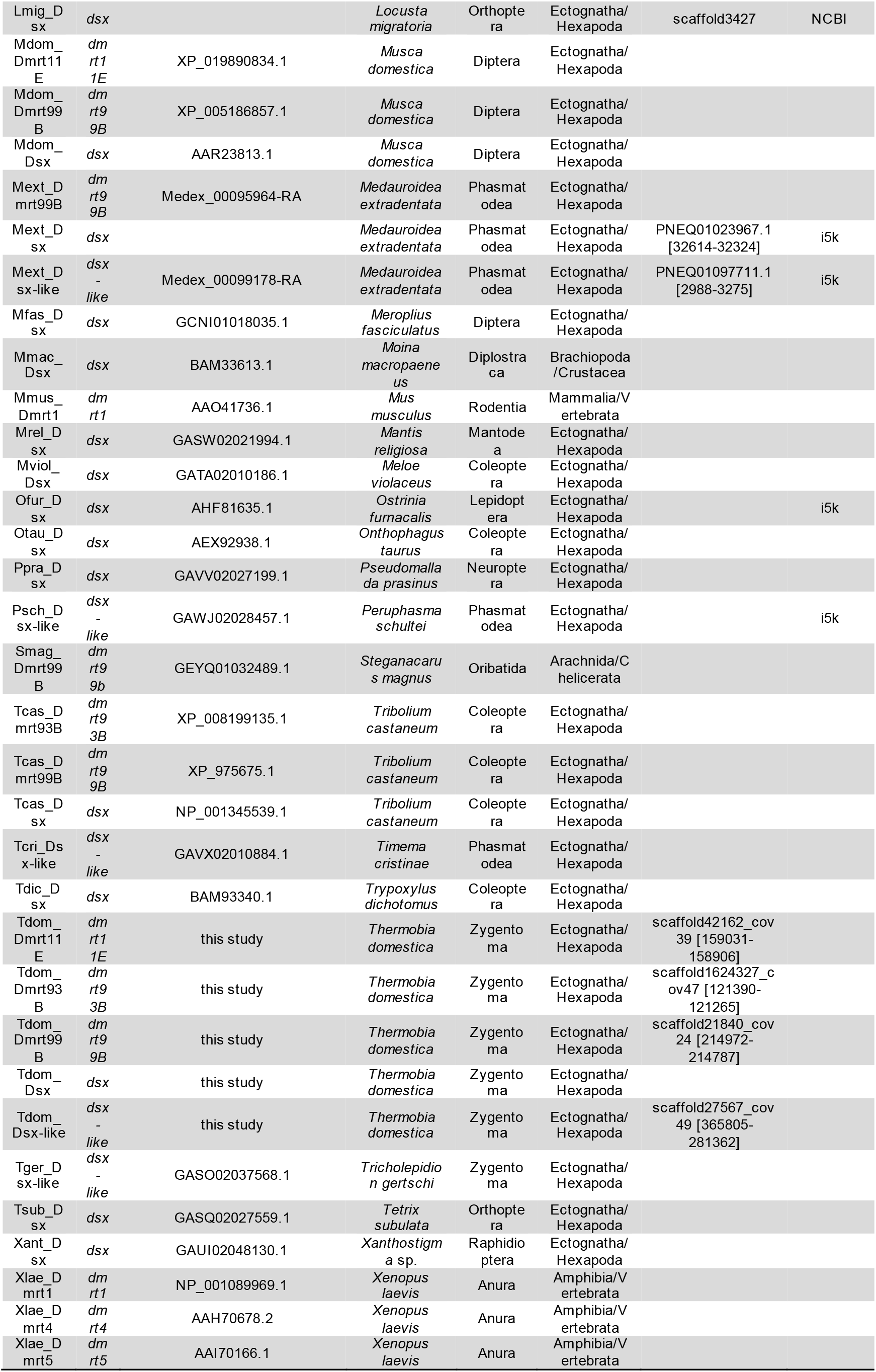
Taxa and proteins used for molecular phylogenetic analysis of DMRT family.

**Table 2.**
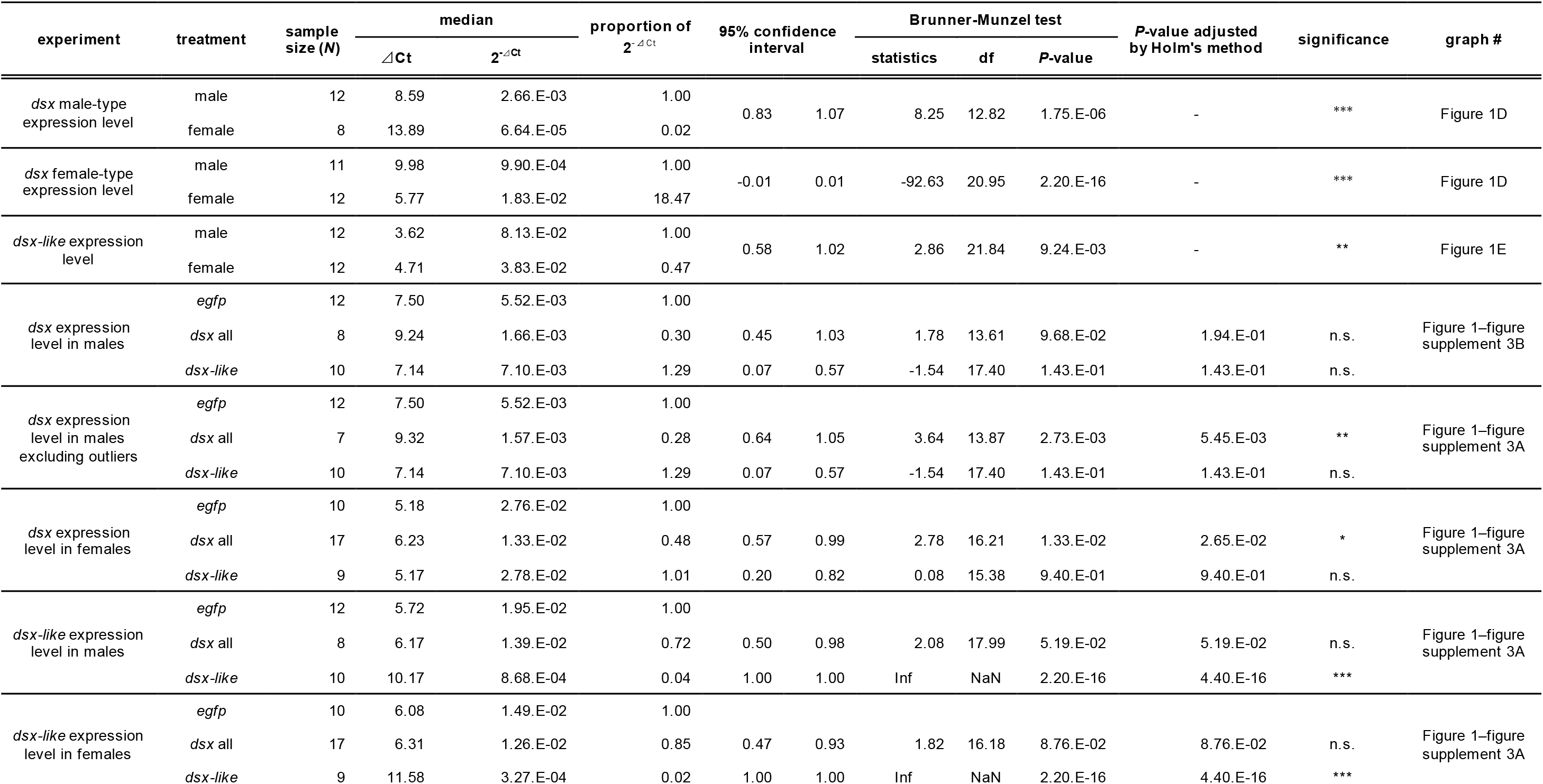
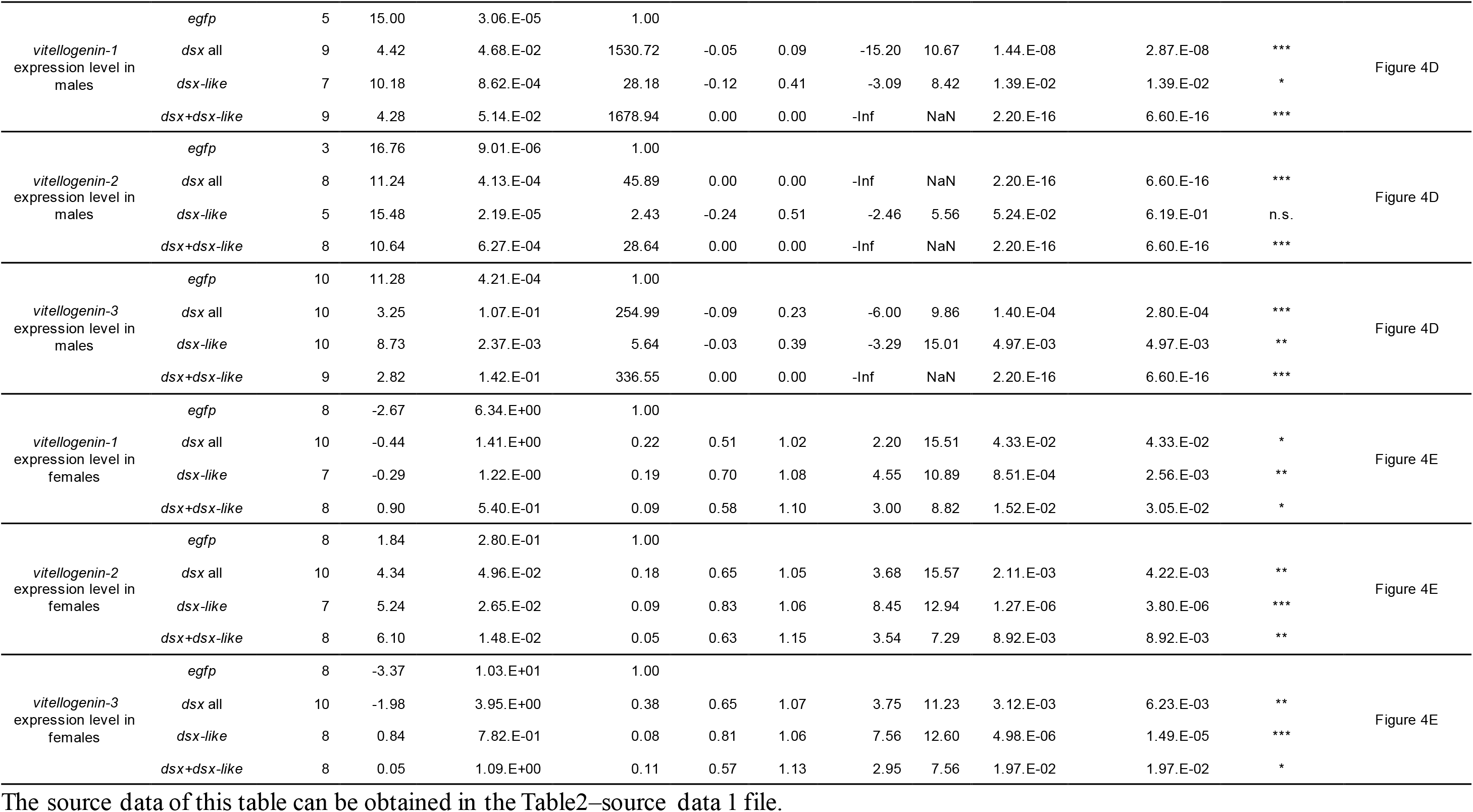
Results of RT-qPCR assay and Brunner–Munzel test. Significant levels are indicated by asterisks in significance column: \**P* < 0.05, \*\**P* < 0.01, \*\*\**P* < 0.001. n.s. means non-significance.

### Sex-specific splicing regulation of *doublesex* in *Thermobia domestica*

We detected two major isoforms of *dsx* of *T. domestica* (Figure 1C). Mapping these sequences to our genome data showed that the longer (951 bp) and shorter (756 bp) isoforms. Both isoforms shared a Doublesex and Mab-3 (DM) domain-containing exon, but differed in the 3’-terminus. The longer isoform was expressed at 40-fold higher levels in males than in females in the fat body. The shorter isoform was expressed in the fat body 18 times higher in females than in males (Figure 1D). We named thus the longer and shorter isoforms *dsx* male-type and *dsx* female-type, respectively. *dsx-like* was expressed two-fold more highly in males than in females (Figure 1F) but had no isoform (Figure 1E). Thus, *dsx* was regulated by a sex-specific splicing, whereas *dsx-like* was not under splicing control.

Before RNA interference (RNAi) analyses, we confirmed that expression levels of *dsx* and *dsx-like* mRNA were significantly reduced in *dsx* and *dsx-like* RNAi males and females. (Figure 1–figure supplement 3A and Table 2; see Materials and Methods section).

### Single-functionality of *doublesex* for phenotypic sex differentiation in *T. domestica*

The sexual traits in *T. domestica* are the male penis and the female ovipositor (Figure 2A–D).

**Figure 2.**
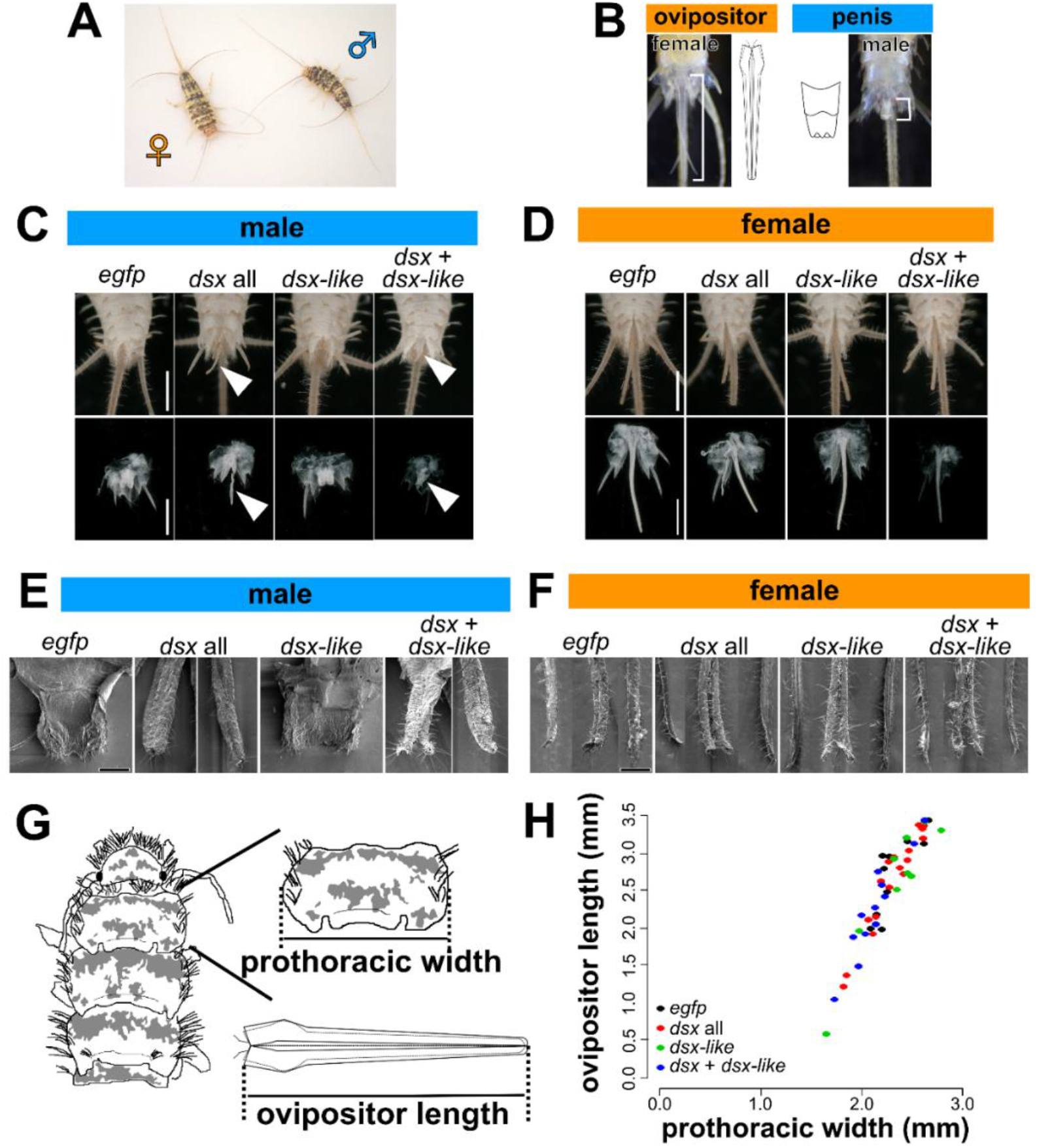
Function of Doublesex in sexually dimorphic traits in *Thermobia domestica*. (A) A pair of *T. domestica*. The female looks much the same as the male. (B) Sexually dimorphic traits of *T. domestica*. Females possess an ovipositor and males have a penis. (C to F) Phenotypes of *dsx* and *dsx-like* RNA interference (RNAi) individuals. Light microscopic image (C and D) and scanning electron microscopic image (E and F). Scales: 1 cm (C and D); 50 µm (E and F). (G to H) Effect of *dsx* and *dsx-like* RNAi on the growth of the ovipositor. The ovipositor length was measured and is plotted with prothoracic width, an indicator of body size. The measurement scheme (G) and scatter plots (H). Each plot in (H) indicates a value of each individual. Sample size of (H) is listed in Table 4. Results of the generalized linear model analysis are listed in Table 4.

In *T. domestica* males, the penis is an unpaired appendix on the abdomen segment IX (*Matsuda, 1976*). The penis was sub-segmented into two parts. There were many setae on the left and the right side of the distal tips (Figure 2E, Figure2–figure supplement 1E). The surface of the penis had a reticulated pattern (Figure 2E, Figure2–figure supplement 1C, E). This simple penial structure was presumably gained at the last common ancestor of Ectognatha (*Boudinot, 2018*).

In *T. domestica* female, the ovipositor consists of two pairs of appendices (gonapophysis) and is derived from the retracted vesicles on the abdomen VIII and IX (*Emeljanov, 2014*; *Matsuda, 1976*). This ovipositor is an autapomorphy of Ectognatha (*Beutel, 2017*; *Kristensen, 1975*). The gonapophyses on the abdomen VIII (valvula I) were the ventral part of the ovipositor and a paired structure. The gonapophyses on the abdomen IX (valvula II) were the dorsal side of the ovipositor and were united to form an unpaired structure (Figure 2–figure supplement 2B). The distal tip of the valvula II remained a paired structure and possessed dense setae (Figure 2–figure supplement 2A). The valvula I was shorter than the valvula II (Figure 2–figure supplement 2A). Both valvulae were sub-segmented and have some setae (Figure 2F). The valvula I and II were connected through a tongue-and-groove structure (olistheter). The olistheter consisted of an aulax (”groove”) on the valvula I and a rhachis (”tongue”) on the valvula II (Figure 2–figure supplement 2B). Within the valvulae, the epithelial cells were beneath the cuticular layer. The cuticular layer was thickened and multi-layered in the outer surface of the ovipositor. In contrast, the inner surface (i.e., the side of the egg cavity) of the ovipositor had a thin and single-layered cuticle.

In *dsx* or *dsx* male-type RNAi males, a tubular organ was formed instead of the penis (Figure 2C, Figure2–figure supplement 1A). This tubular organ consisted of two pairs of appendage-like structures. The inner one is connected to the gonopore and the ejaculatory duct. The outer one had a lot of setae on its tip (Figure 2E, Figure2–figure supplement 1C, E). Thus, the inner pair was similar to the valvula I of the female ovipositor and the outer one was similar to the valvula II. These features indicated that the tubular organ in the *dsx*RNAi males was parallel to the female ovipositor. The same phenotype was found in the *dsx* and *dsx-like* double RNAi males. In contrast, the *dsx-like* males possessed a penis the same as that of the control insects.

In females that were treated with RNAi for *dsx*, *dsx* isoforms, *dsx-like*, or both genes, the external genital organ was the same as the ovipositor of the control females that described in the above section (Figure 2D, F, Figure 2–figure supplement 1B, D, F, Figure2–figure supplement 2). Thus, in the view of histology, the location, and the relation to other elements, the external organ of the RNAi females was not different from the ovipositor of the control ones.

### No evidence for the effects of *doublesex* on female phenotypes in *T. domestica*

To investigate whether *dsx* in *T. domestica* females has functions other than female differentiation for phenotypes, we examined its role in the growth and maintenance of female organs (Figure 2G). We did not detect a significant difference in ovipositor length and prothoracic width, an indicator of body size, between the controls and the *dsx* and *dsx-like* RNA females (Figures 2H, Figure 2–figure supplement 3, Figure2–figure supplement 4 and Table 4).

We then examined the effects of *dsx* on gonads, reproductive systems, and gametogenesis in *T. domestica*.

In males of *T. domestica*, a pair of testes was consisted of some testicular follicles (Figure 3A). Each testicular follicle was connected to the vas deferens via the vas efferens (Figure 3–figure supplement 1B). The seminal vesicle lay between the vas deferens and the ejaculatory duct. A pair of the ejaculatory ducts was associated with each other in the front of the gonopore in the penis (Figure 3C). The testicular follicles were a bean-like shape and the seminal vesicles were a bean pod-like shape. In the testicular follicle, the spermatogonia was in the antero-most part (Figures 3A). The primary and secondary spermatocytes lay in the middle part (Figures 3A). In the posterior part of the testicular follicle, there were some sperm bundles (Figures 3A). The wall of the testicular follicle consisted of a single flattened epithelial layer.

**Figure 3.**
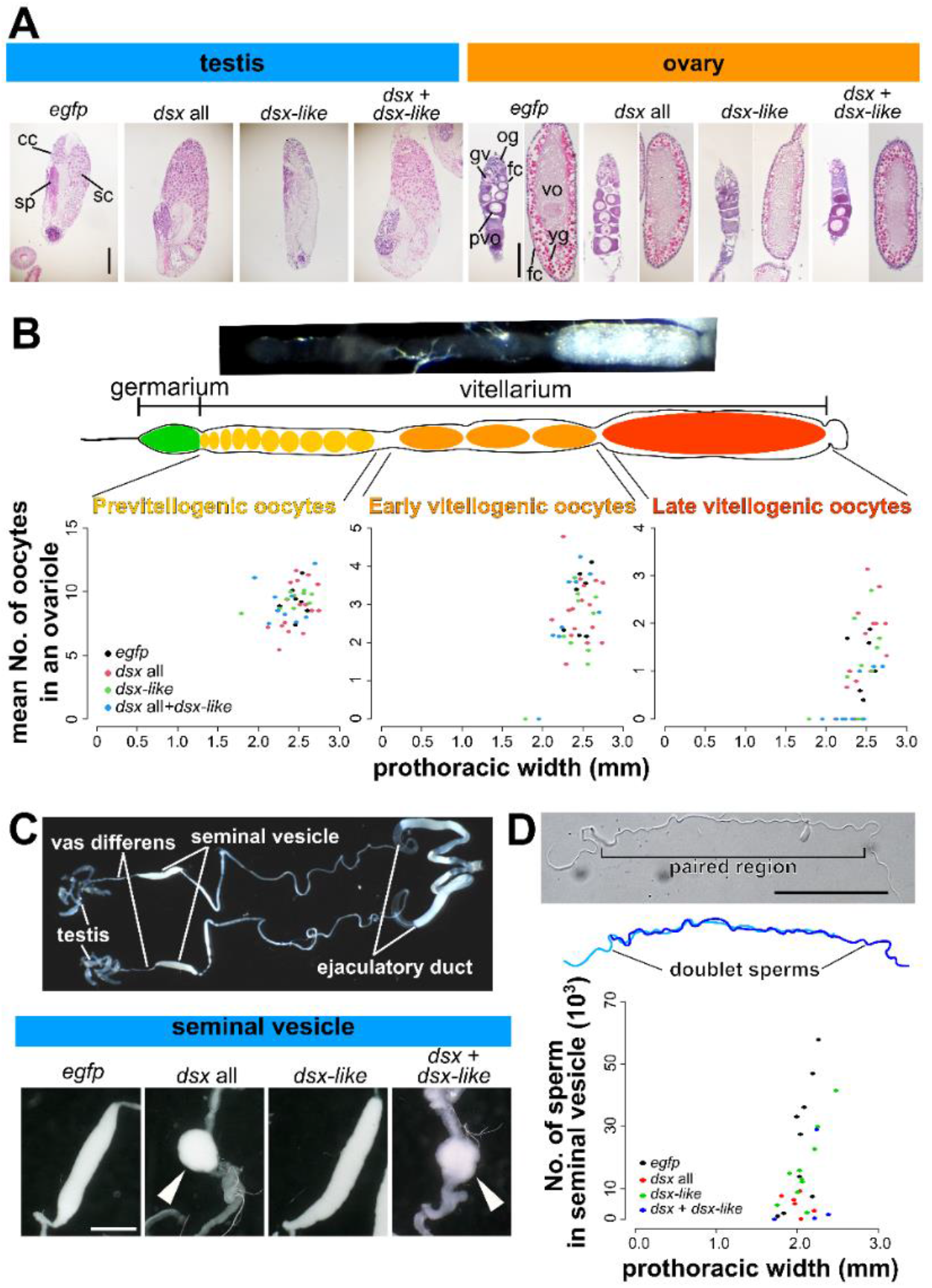
Function of *doublesex* in reproductive systems and fecundity. (A) Effects of *dsx* and *dsx-like* RNAi on gonad morphology and gametogenesis. In images of the ovary, the left and right panel in each treatment show germarium/previtellogenesis and vitellogenesis, respectively. cc, cystocyte; fc, follicle cell; gv, germinal vesicle; og, oogonia; pvo, previtellogenic oocyte; sc, spermatocyte; sp, sperm; yg, yolk granule; vo, vitellogenic oocyte. (B) The effect of *dsx* and *dsx-like* RNAi on oocyte number in each oogenetic stage. The upper panels are images of an ovariole in *T. domestica*. The lower panel shows scatter plots of oocyte numbers and prothoracic width. Results of the generalized linear model analysis are in Tables 4 (female) and 5 (male). (C) The effect of *dsx* and *dsx-like* RNAi on the seminal vesicle. The upper panel is a gross image of the male reproductive system. The lower one shows the phenotype of each treatment. (D) The effect of *dsx* and *dsx-like* RNAi on sperm number in seminal vesicles. The upper panels are images of sperm in *T. domestica*. The sperm of *T. domestica* is usually paired with other sperm. The lower panel shows a scatter plot of sperm numbers by prothoracic width. Scales: 50 µm (A and C); 10 µm (D). Sample size are also listed in Tables 4 (D) and 5 (C).

We observed the above features of the reproductive system in *dsx* or *dsx-like* RNAi males (Figure 3A and C). In *dsx* and both genes RNAi males, the seminal vesicles were round in shape. The vas efferens was filled with the sperm (Figure 3–figure supplement 1B). In contrast, we could not find differences in the morphology of the testicular follicles or spermatogenesis between the RNAi and control males. The male reproductive system and spermatogenesis showed no visible difference between *dsx* female-type and *dsx-like* RNAi females and the control ones.

In females of *T. domestica*, each ovary consists of five ovarioles and was attached to the anterior part of the abdomen via the terminal tuft (Figure 3–figure supplement 1). The ovarioles were associated with each other at the lateral oviduct. The lateral oviduct was connected to the common oviduct and subsequently opened at the gonopore in the valvula I. There was no vagina between the gonopore and the oviduct. The spermatheca was located on the branch point of the common oviduct along the midline (Figure 3–figure supplement 1A). The spermatheca was divided into two parts: anterior and posterior (Figure 3–figure supplement 2). The anterior part consisted of a pseudostratified layer of the columnar epithelial cells that were secretory. The posterior part was surrounded by a single layer of epithelial cells. The ovariole was panoistic-type and was composed of two parts: the germarium and the vitellarium (Figure 3A, B). The germarium contained many oogonia and young oocytes. The vitellarium had previtellogenic and vitellogenic oocytes. The oocytes in the vitellarium were surrounded by a single layer of follicle cells. There were pedicel cells in the terminal of the ovariole. The previtellogenic oocyte had a large germinal vesicle and basophilic cytoplasm. The vitellogenic oocyte was elongated along the anterior-posterior axis of the ovariole and had eosinophilic cytoplasm. Many eosinophilic lipid droplets were present in the peripheral region of the vitellogenic oocytes. The follicle cells were flattened and columnar in shape in the previtellogenesis and the vitellogenesis.

We observed the above features of the reproductive system in *dsx* or *dsx-like* RNAi females (Figures 3A, Figure 3–figure supplement 1, Figure 3–figure supplement 2). We could not detect differences in the female reproductive system or oogenesis between the RNAi females and the controls. This result suggests that the *dsx* and *dsx-like* have no function in the formation of female traits and gametogenesis at the tissue and cellular level.

Also, we were not able to detect any differences in the oocyte number and size between the controls and RNAi females (Figures 3B, Figure3–figure supplement 3 and Table 4). However, the seminal vesicle in males became round in shape in the *dsx* RNAi males in contrast to the bean pod-like shape observed in the control males (Figures 3C, Figure3–figure supplement 1). We detected a significant reduction of the number of sperms within the seminal vesicle in *dsx* RNAi males (Figure 3D and Table 5). *dsx* thus contributed to the development of the reproductive system and gametes in males, but not in females, of *T. domestica*.

Our results cannot show evidence for roles of *dsx* in *T. domestica* females.

### Cryptic role of *doublesex* for female-specific transcripts in *T. domestica*

Previous studies in Hemimetabola (*Takahashi et al., 2021*; *Wexler et al., 2015*; *Zhuo et al., 2018*), and our results in Zygentoma show that *dsx* is not essential for the formation of female phenotypes in non-Holometabola. In contrast, considering the fact that the *dsx* female-specific isoform retains ORFs in non-Holometabola, there is a still possibility that *dsx* may have some function in non-holometabolan females. To investigate this functionality, we focused on the effect of *dsx* on female-specific gene expression.

We focused on the *vitellogenin* (*vtg*) gene, one of the major egg yolk proteins, which is specifically expressed in females of Bilateria (Byrne et al., 1989; *Hayward et al., 2010*). In the Holometabola, *dsx* promotes *vtg* mRNA expression in females and represses it in males (*Shukla and Palli, 2012*; *Suzuki et al., 2003*; *Thongsaiklaing et al., 2018*). Our RNA-seq analysis showed that levels of the three *vtg* mRNAs in *T. domestica* were more than 10000-fold higher in females than in males (Figure 4A–C). We found that the *vtg* mRNAs were expressed at 40–100-fold higher levels in *dsx* RNAi males than in the control males (Figure 4D and Table 2). Notably, the *dsx* RNAi females produced around half the amount of *vtg* mRNA as the controls. The *dsx-like* RNAi and a double-knockdown of *dsx* and *dsx-like* also significantly reduced the expression of *vtg* in females (Figure 4E and Table 2).

**Figure 4.**
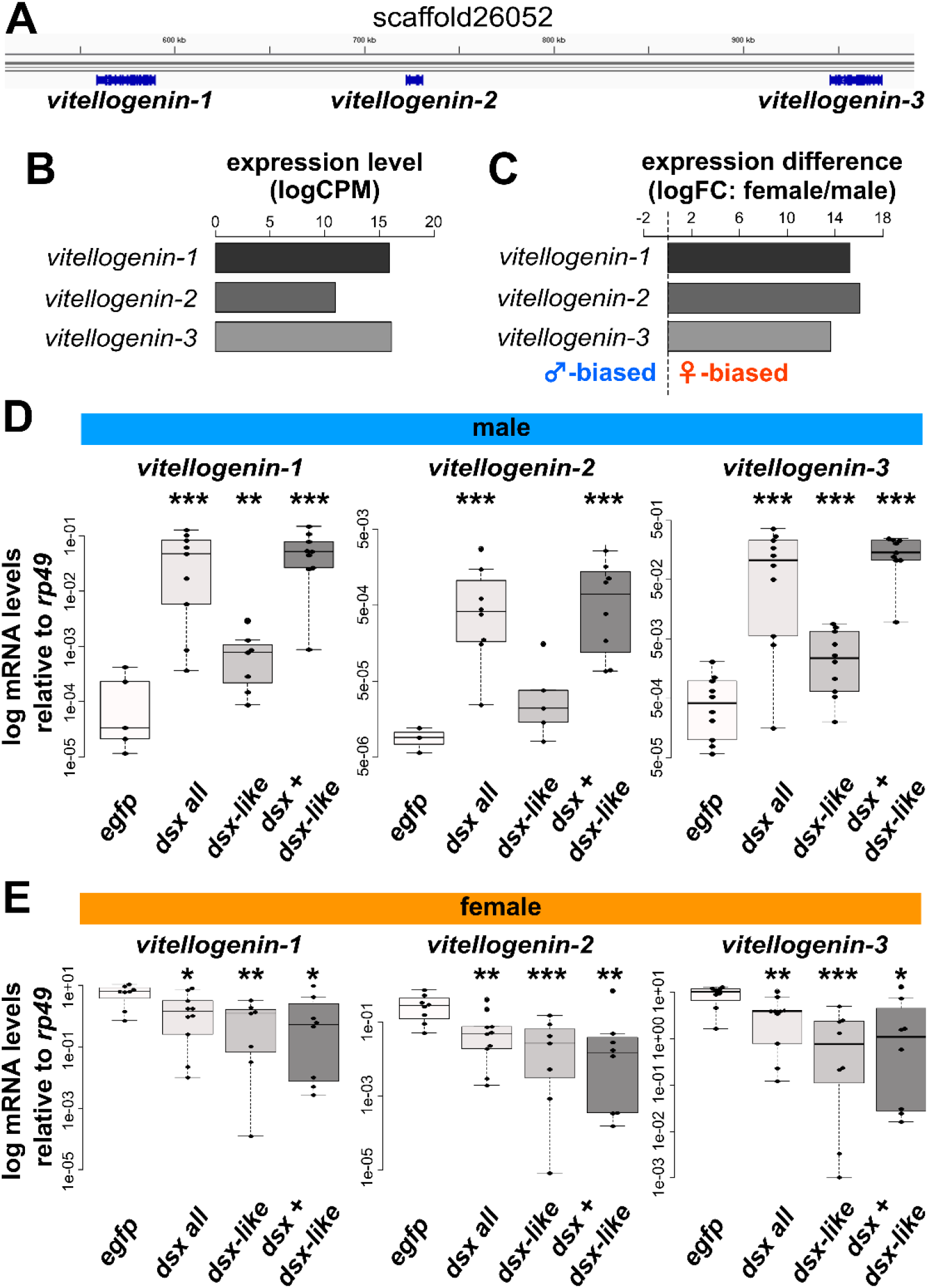
Function of *doublesex* in *vitellogenin* gene expression. (A to C) Features of *vitellogenin* (*vtg*) genes of *Thermobia domestica*. An image of the *vtg* gene locus in an Integrated Genome Viewer (A). The *vtg* genes are highly expressed in females (B and C). The source data of B and C are provided as Figure4–source data 1. (D and E) Effect of *dsx* and *dsx-like* RNAi on *vtg*genes’ expression in males (D) and females (E). The decreases were about 2/5–1/5, 1/10–3/50, and 1/9–1/20 in *dsx*, *dsx-like*, and both the genes RNAi females. The mRNA expression levels are shown as logarithmic scales. Each plot represents an individual. Sample sizes are listed in Table 2 (C and D). Results of Brunner–Munzel tests are indicated by asterisks: \**P* < 0.05; \*\**P* < 0.01; \*\*\**P* < 0.001 and is also described in Table 2. *P* > 0.05 is not shown.

This result is the first case that *dsx* can promote female-specific *vitellogenin* expression in non-holometabolan species, even though it does not affect female phenotypes such as the oocyte size and number. This finding suggests that it has opposite functionality between males and females in *T. domestica* at the gene-regulatory level.

### Differences of *doublesex* sequences between single- and dual-functional species

Genes can gain new functions due to gene duplication, co-factor function, changes in *cis*- or *trans*-region (*Carroll, 2005*; Ganfornina and Sánchez, 1999; *Mann and Carroll, 2002*), or acquiring new exons. *dsx* paralog (*dsx-like*) found in this study did not contribute to female phenotypes (Figure 2 and 3). The female and male isoforms of *dsx* share the same DNA-binding domain, and *intersex*, a co-factor gene of *dsx*, contributes to the formation of female traits in *Nilaparvata lugens* (*Zhang et al., 2021*), which has the single-functional *dsx*. These facts indicate that the gene duplication, neo-functionalization of co-factors, and changes in *cis*-regulatory elements are not likely to contribute to the evolution of the dual-functionality. Thus, we explored the remaining possibilities: novel exons or *trans*-regions.

We found that there are alternative splice types in *T. domestica*, which has an alternative 5ʹ splice site, and in Pterygote insects, which have a mutually exclusive exon, but did not find any differences of exon structure between species with single-functionality of *dsx*, and those with dual-functionality (Figure 5A).

**Figure 5.**
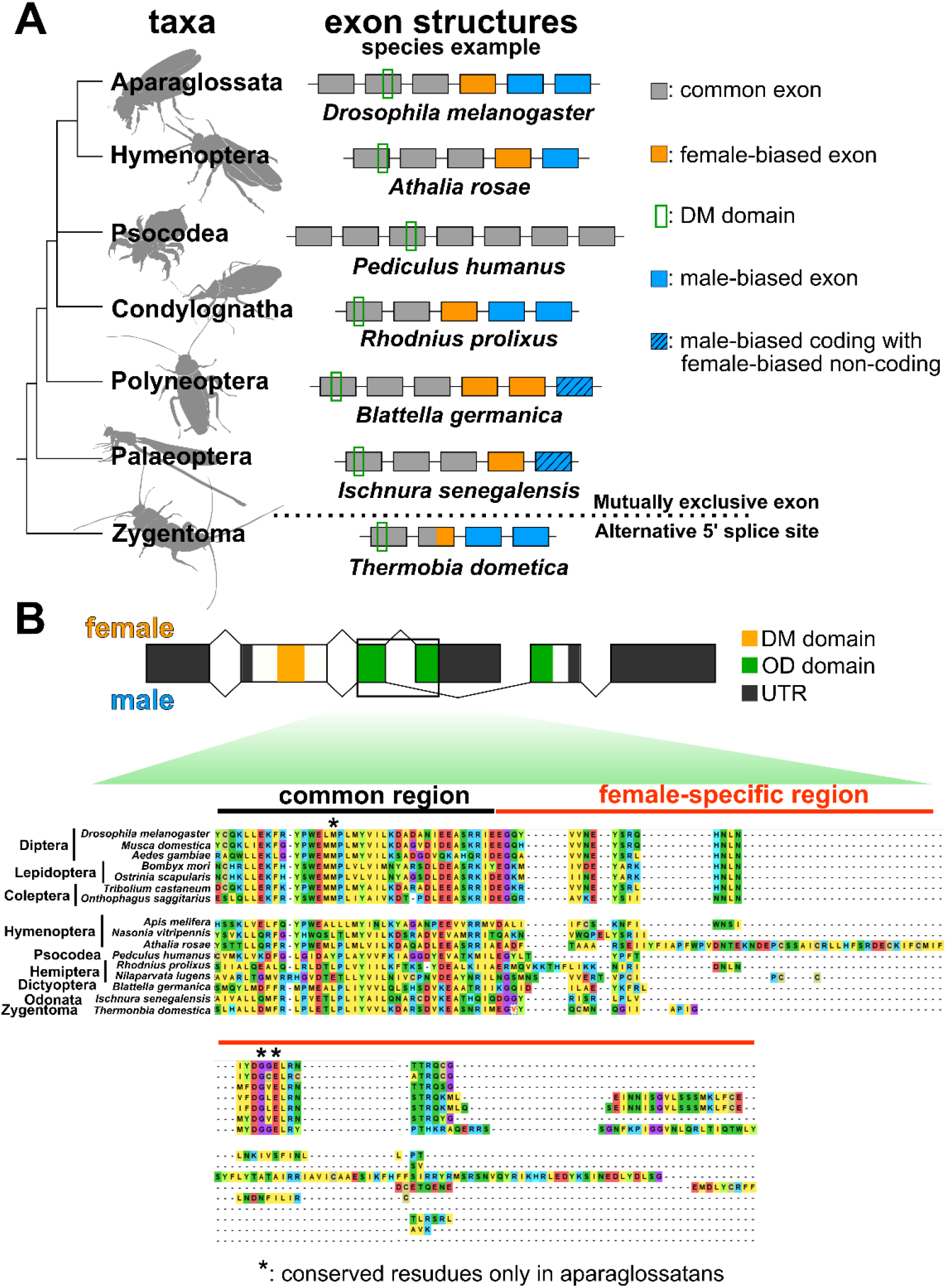
Comparisons of molecular features in *doublesex* among insect taxa. (A) Exon structure of *dsx* among insect taxa. The coding region of *dsx* is shown. The phylogenetic relationship is that of *Misof et al. (2014)* (B) amino acid sequence of Dsx among insect taxa. The upper image shows the *dsx* structure of *Drosophila melanogaster*. Asterisks indicate conserved residues in Aparaglossata.

We finally discovered amino acid sequences in the female-specific region that differed between taxa with single- and dual-functionality of *dsx*. *dsx* isoforms are sexually different in the OD domain. This domain was divided into common, female-specific, and male-specific regions. Our multiple alignment analysis revealed that the female-specific region was highly conserved among the taxa with dual-functionality of *dsx*, but not among those with single functionality (Figures 5B, Figure 5–figure supplement 1). In contrast, the male-specific region was not highly conserved among the taxa with dual-functional *dsx* (Figure 5–figure supplement 2).

## Discussion

### A novel type of the output in the sex differentiation system

We show that the *dsx* of *Thermobia domestica* is essential for producing male phenotypes, but does not contribute to female phenotypes. In contrast to the phenotypes, this study showed that the female-specific *vitellogenin* genes are slightly promoted by *dsx* in *T. domestica* females. These facts indicate that *dsx* in *T. domestica* has an opposite transcription-regulatory function in males and females. Therefore, in *T. domestica*, *dsx* contributes only to male differentiation at the phenotypic level, but affects both sexes at the transcription-regulatory level (seemingly useless nature).

There have been two known types of outputs of the insect sex differentiation system: one that can regulate feminization via both transcription-regulation and phenotypes, and one that cannot. The former is found in Diptera, Lepidoptera, and Coleoptera (e.g., Luo and Baker, 2015; Suzuki et al., 2003; Shukla and Palli, 2012), the latter in Dictyoptera and Hemimetabola (*Wexler et al., 2019; Zhuo et al., 2018*). In some species, such as the dung beetle *Onthophagus taurus*, *dsx* contributes only to male trait formation in some tissues (*Ledón-Rettig et al., 2017*). In these species, *dsx* is also responsible for producing traits of both sexes in other tissues. Therefore, it is likely that the former type is the primary capability of the sex differentiation systems in these species and that the function for promoting female differentiation may tissue-specifically become silent. In contrast, the seemingly useless nature of *dsx* for females in *T. domestica* is a third type of the insect sex differentiation system (Figure 6A). This third type indicates that phenotypic and transcription-regulatory levels do not necessarily coincide in the output of sex differentiation systems.

**Figure 6.**
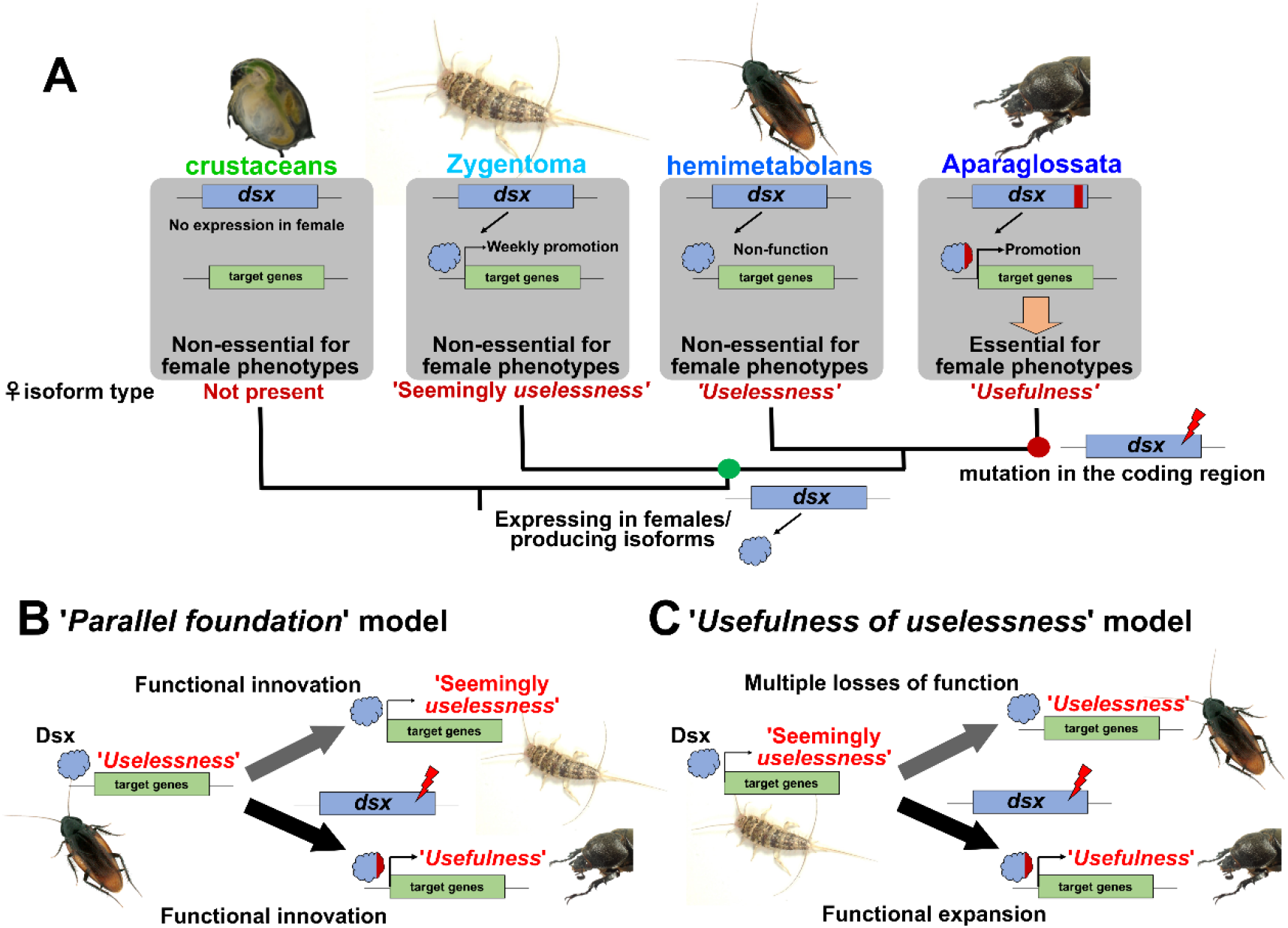
Evolutionary transition of outputs of the insect sex differentiation system. (A) phylogenetic mapping of functionality of *dsx* for females. crustaceans: *Kato et al. (2011)*; Zygentoma: this study; hemimetabolans: *Wexler et al. (2019), Zhuo et al. (2018);* Aparaglossata: e.g., *Shukla and Palli (2012), Suzuki et al. (2003)*. (B) ‘*Parallel foundation*’ model. (C) ‘*Usefulness of useless*’ model.

The sex differentiation systems of crustaceans, vertebrates, and nematodes have DMRT family transcription factors as bottom factors that are responsible for promoting male differentiation (*Kato et al., 2011*; *Kopp, 2012*; *Raymond et al., 1998*; *Raymond et al., 2000*). No hierarchical discrepancy has been found in the output of the sex differentiation system of these animals. Therefore, the output of the sex differentiation system in most animals can be classified into three categories: that are capable of contributing to 1) only masculinization (crustaceans, nematodes, vertebrates, Hemiptera, and Dictyoptera), 2) both masculinization and feminization (Diptera, Lepidoptera, and Coleoptera) or 3) both masculinization and feminization at the transcription-regulatory level but only masculinization at the phenotypic level (*T. domestica*).

### On the origin of the sex-specific splicing of *doublesex*

In pterygote insects (e.g., *Burtis and Baker, 1989*; *Shukla and Palli, 2012*; *Mine et al., 2017*; *Ohbayashi et al., 2001*; *Takahashi et al., 2019*; *Wexler et al., 2019*; *Zhuo et al., 2018*), *dsx* has sex-specific isoforms, except for a termite *Reticulitermes speratus* (*Miyazaki et al., 2021*). In contrast, *dsx* is controlled via a male-specific transcription in chelicerates and crustaceans (*Kato et al., 2011*; Li et al., 2018; *Panara et al., 2019*; *Pomerantz et al., 2015*). Here, sex-specific splicing regulation is observed in *dsx* of *T. domestica*, suggesting that sex-specific splicing regulation originated between the common ancestor of Branchiopoda and Hexapods and the common ancestor of Dicondylia (Zygentoma + Pterygota) (Figure 6A).

### On the origin of the role of the insect sex differentiation system for females

There are discrepancies in the output of the sex differentiation system in *T. domestica* at the phenotypic and transcription-regulatory levels. In such a case, it has been proposed to map the characteristics of each hierarchical level separately on a phylogenetic tree (c.f., *Abouheif, 1999*).

At the phenotypic level, previous studies have focused on taxon-specific or highly complex sex differences, leaving open the possibility of a secondary loss of function of *dsx* for females. However, the single-functionality of *dsx* even in *T. domestica*, which has an ancestral, simple sex difference, supports a single origin for *dsx* single-functionality. Based on a recent phylogenetic hypothesis (*Beutel et al., 2017*; *Misof et al., 2014*), the transition from the single-to dual-functionality is estimated to have occurred in the common ancestor of Aparaglossata (holometabolan insects excluding Hymenoptera) (Figure 6A). At the phenotypic level, our findings ensure the evolutionary model by *Wexler et al. (2019)* that *dsx* initially acquired sex-specific isoforms and later became essential for female differentiation.

At the level of gene regulation, *dsx* in *T. domestica* can slightly promote the transcription of female-specific genes. Therefore, it is estimated that the common ancestor of Dicondylia already possessed the sexual dimorphic transcriptional control of *dsx*. In this case, it is inferred that the transcription-promoting function of female-specific genes was secondarily lost in hemimetabolan insects. Alternatively, it is possible that the common ancestor of the Dicondylia had also the single-functionality at the level of transcriptional regulation and that Zygentoma and Aparaglossata independently acquired the ability to promote transcription in females.

The question is how the function of *dsx* for the female phenotype evolved. We have found differences in the sequence of the female-specific region of *dsx* between phenotypically single-functional and dual-functional taxa. This region is located in the oligomerization (OD) domain, interacts with transcription factors (*An et al., 1995*; *Erdman et al., 1996*; *Ghosh et al., 2019*; *Romero-Pozuelo et al., 2019*) and transcriptional mediators such as *intersex* which is essential for female differentiation (*Gotoh et al., 2016*; *Morita et al., 2019*; *Xu et al., 2019*; *Yang et al., 2008*). Studies in *Blattella germanica* described low conservation of the OD domain (*Wexler et al., 2019*). *Baral et al. (2019)* also reported that the rate of non-synonymous substitutions in the female-specific region is low in Aparaglossata and high in Hymenoptera. However, the evolutionary significance of this region has been unclear. Our result suggests that the accumulation of mutations in female-specific regions has led to the female-differentiating functions at the phenotypic level (Figure 6). It was theoretically predicted that non-functional isoforms gain functions through mutation accumulation in coding regions (*Keren et al., 2010*). The functional evolution of *dsx* in insects may fit this prediction.

### Evolutionary scenarios for the transition of the output of the sex differentiation system

In conclusion, we propose two alternative scenarios for the transition of the outputs of the insect sex differentiation system. “*Parallel foundation*” hypothesis (Figure 6B): *dsx* has independently acquired the function for promoting female transcription in Zygentoma and Aparaglossata, and also gained the entirely novel role in phenotypic differentiation in females through the accumulation of coding mutations in Aparaglossata. This is the hypothesis that a useful functionality has arisen from a completely useless functionality. “*Usefulness of uselessness*” hypothesis (Figure 6C): *dsx*, through the accumulation of mutations in its coding region, enhanced its weak transcription-promoting ability in females and became essential for producing the female phenotypes. This is the hypothesis that a useful functionality arose from a seemingly useless functionality.

Both these hypotheses can explain the diversity in outputs of the sex differentiation system via the accumulation of coding mutations in existing genes. Thus, a heterotypic evolution may drive the evolution of outputs of the sex differentiation systems. Textbook examples of the heterotypic evolution include a mutation of human *hemoglobin β* leading to sickle cells and coding mutations of *Ultrabithorax* that resulted in suppressed leg development in insects (reviewed in *Arthur, 2010*; *Gilbert, 2013*; *Futuyma and Kirkpatrick, 2017*). So far, heterotypy at the molecular level is considered as the change in the molecular function leading to entirely novel or no functions from existing functions of the molecule in phenotypes. On the other hand, the female isoform of *dsx* shows no function in female phenotypes. Our hypotheses are models that a non-functional isoform gains functionality at the phenotypic level. The heterotypic evolution may need to be divided into modification from existing functions and innovation from non-functionality at phenotypic level.

In this study, we succeeded in detecting the transcription-promoting ability of *dsx* in females by examining the expression of *vitellogenin*. In hemimetabolan insects. the transcription-promoting function in females has not been found in *vitellogenin* (*Wexler et al., 2019*; *Zhuo et al., 2018*), but might be detected in other female-specific genes. Future comprehensive examination of the effects of *dsx* by transcriptome and comparisons at the mega-evolutionary level, i.e., higher taxa than families (*Arthur, 2003*), using broad insect taxa will test these hypotheses.

## Materials and Methods

### Animals

A firebrat, *Thermobia domestica* (Packard, 1873), was used as an emerging model apterygote. *T. domestica* is one of the species belonging to the Zygentoma (Lepismatidae). The insects were kept at 37°C in total darkness condition and fed with fish food (TetraFin Goldfish Flakes, Tetra GmbH, Melle, Germany) in our laboratory. Stock colonies were reared in plastic cases of 30 cm×40 cm or 18 cm × 25 cm in length. Eggs were collected from tissue paper in the case and incubated at 37°C. For nymphal RNAi analysis, colonies of hatched nymphs were reared up to the fourth instar in a six-well plate and then transferred into 24-well plates to be kept individually. For adult RNAi analysis, adult insects were collected from the stock colony and transferred into the plates. For nymphal RNAi analysis, we used firebrats from April to June, 2019, February to April, April to July, and September to December, 2020. For adult RNAi, adult firebrats were manipulated from June to July, 2020.

### Estimation of molt timing

Estimating the molt timing of insects is essential for the analysis of developmental processes and the functions of developmental regulatory genes. The timing of Hemi- or holometabolan insects can be estimated using morphological changes such as a wing growth. However, timing is hard to estimate in apterygote insects since they have little change in their morphology during postembryonic development. *T. domestica* forms scales in the fourth instar, and changes the number and length of its styli during the fourth to ninth instar under our breeding conditions. These features can be used to estimate molt timing, but it is difficult to apply these criteria to experiments using adults or a large number of nymphs. To resolve this problem, we used leg regeneration after autotomy and time-lapse imaging to estimate the molt timing of *T. domestica*. Autotomy occurs at the joint between the trochanter and femur in *T. domestica*. An autotomized leg regenerates after one molt (*Buck and Edwards, 1990*). For nymphal RNAi analysis, we amputated a right hindleg at the autotomic rift, using tweezers, and observed whether the leg had regenerated. This test enabled us to rapidly estimate the molt timing. For adult RNAi, time-lapse imaging was used to determine the precise time of molt. We build a time-lapse imaging system with a network camera system (SANYO, Tokyo, Japan) set in an incubator at 37°C (Figure 4–figure supplement 1A). Photos of insects in the 24-well plate were taken every five minutes. We created a time-lapse movie from the photos every 12 hours using ImageJ 1.52a (https://imagej.nih.gov/ij/) and observed whether the insects molted (Figure 4–figure supplement 1B, Video 1).

### De novo genome assembly

A whole genome of *T. domestica* was sequenced to analyze the exon-intron structure of *dsx*. We selected an adult female of *T. domestica* from our stock colony and removed its alimentary canal. Genomic DNA was extracted from the sample using DNeasy Blood and Tissue Kit (QIAGEN K.K., Tokyo, Japan). A paired-end library was constructed from 1 µg of the DNA using TruSeq DNA PCR-Free LT Sample Prep kits (Illumina K.K., Tokyo, Japan) following the manufacturer’s instructions. The library was run on a sequencer (HiSeq 2500; Illumina K.K., Tokyo, Japan). We obtained 417 Gb of raw reads and assembled them using Platanus v1.2.4 assembler (*Kajitani et al., 2014*) after removal of the adapter sequences.

### Transcriptome analysis

To search for *doublesex* (*dsx*) and *vitellogenin* (*vtg*) homologs, we performed RNA-seq analysis. Adults of 15 ♀♀ and 15 ♂♂ of *T. domestica* were sampled 1440 minutes after a molt in December, 2019. The fat bodies of the individuals were removed using tweezers in a phosphated buffered saline (PBS; pH=7.2). Three adults were used per sample. Total RNA was extracted from 10 samples (5♀♀, 5♂♂) using RNeasy Micro kits (QIAGEN K.K., Tokyo, Japan) following the manufacturer’s instructions. The concentration of purified RNA was measured using a Qubit 4 fluorometer (QIAGEN K.K., Tokyo, Japan) with Qubit RNA BR Assay kits (QIAGEN K.K., Tokyo, Japan). Paired-end libraries were constructed from 100 ng of the total RNAs using TruSeq RNA Library Prep kits v2 (Illumina K.K., Tokyo, Japan) following the manufacturer’s instructions. The libraries were run on a sequence (Hiseq, Illumina, Tokyo, Japan). The library preparation and sequencing were performed by Genewiz Strand-Specific RNA-seq service. We mapped the reads obtained to the assembled genome using the HISAT2 program (Kim et al., 2019) with a default option and counted the mapped reads using the STRINGTie program (Pertea, 2015) with default parameter settings. Differential expression gene analysis was performed based on the count matrix using the “edgeR” package *(Robinson et al., 2010*) in R-v4.0.3 (*R Core Team, 2020*). Information about the samples can be obtained from the National Center for Biotechnology Information (NCBI) BioSample database (Accession number: SAMN18175012–SAMN18175021).

### Molecular phylogenetic analysis

Dsx is a member of the Doublesex and Mab-3 Related transcriptional factors (DMRT) family, and has a DNA binding domain, Doublesex and Mab-3 (DM) domain. Pancrustacea generally has four DMRT family genes, Dsx, Dmrt11, Dmrt93B, and Dmrt99B (*Mawaribuchi et al., 2019*). Phylogenetic analysis of Dsx homologs was performed using the amino acid sequences of the DM domain. We used the Dsx sequences of *D. melanogaster* as a query and obtained 97 metazoan DMRT family proteins from the NCBI and the i5k databases (https://i5k.nal.usda.gov/) and our genome data of *T. domestica* by the BLAST analysis (listed in Table 1). We then aligned the sequences using MAFFT version 7 (*Katoh et al., 2013*) with the -linsi option (to use an accuracy option, L-INS-i) and manually extracted the DM domain, which consisted of 61 amino acids (Figure 1–figure supplement 1). Molecular phylogenetic analysis of the aligned sequences was performed using a maximum likelihood method after selecting a substitution model (JTT matrix-based model) with MEGA X (*Kumar et al., 2018*). Bootstrap values were calculated after 1000 replications.

### Full-length cDNA and exon-intron structures

To elucidate the exon-intron structures of Dsx and Dsx-like, we determined the full-length cDNA sequences using a Rapid Amplification of cDNA Ends (RACE) method and performed a BLAST analysis for our genome database of *T. domestica*. We extracted total RNA from eggs, whole bodies, fat body, and gonads of nymphs and adult females and males of *T. domestica* using TRI Reagent (Molecular Research Center Inc., Ohio, USA) following the manufacturer’s instructions. The total RNAs were treated with RNase-Free DNase I (New England BioLabs Japan Inc., Tokyo, Japan) to exclude remaining genomic DNA and purified by phenol/chloroform extraction and ethanol precipitation. For 5ʹ -RACE analysis, mRNAs were purified from 75 µg of the total RNAs using Dynabeads mRNA Purification kit (Thermo Fisher Scientific K.K., Tokyo, Japan) following the manufacturer’s instruction. We then ligated an RNA oligo at the 5’-end of the mRNA using GeneRacer Advanced RACE kits (Thermo Fisher Scientific K.K., Tokyo, Japan). For 3ʹ-RACE analysis, we ligated an RNA oligo of the SMART RACE cDNA Amplification Kit (Takara Bio Inc., Shiga, Japan) at 3ʹ-end of the total RNA during reverse transcription. First stranded (fs-) cDNA was generated from the RNAs using SuperScript III Reverse Transcriptase (Thermo Fisher Scientific K.K., Tokyo, Japan). We used primers specific to the RNA oligos and performed RACE analysis by nested RT-PCR using Q5 High-Fidelity DNA polymerase (New England BioLabs Japan Inc., Tokyo, Japan). The primers specific to *dsx* and *dsx-like* were made from sequences of the relevant genomic regions and are listed in Table 6. The amplicons were separated using the agarose gel-electrophoresis and cloned using TOPO TA Cloning Kit for Sequencing (Thermo Fisher Scientific K.K., Tokyo, Japan) following the manufacture’s protocol. We used a DH5α *Escherichia coli* strain (TOYOBO CO., LTD., Osaka, Japan) as the host cell. Plasmids were extracted using the alkaline lysis and purified by phenol-chloroform and ethanol precipitation. The nucleotide sequences of the cloned amplicons were determined from the purified plasmids by the Sanger Sequencing service of FASMAC Co. Ltd. (Kanagawa, Japan). We then searched the genomic region of the full-length cDNA sequences of *dsx* and *dsx-like* via local blastn analysis.

### Reverse transcription-quantitative PCR (RT-qPCR)

To quantitative mRNA expression levels, we performed RT-qPCR analysis. For investigating the sex-specific expression profile of *dsx* and *dsx-like*, we used the fat body of adults of *T. domestica* since the sexes can be distinguishable by the external morphology at this stage. Fat bodies also exhibit sex-specific physiological functions in adults. Thirteenth instar individuals and adults after molting were sampled for nymphal and adult RNAi analyses, respectively. The sample sizes are reported in the Table 2. We dissected the individuals in PBS and collected their fat body in 2 ml tubes containing TRI Reagent (Molecular Research Center Inc., Ohio, USA). The fat bodies then were disrupted using a TissueLyser LT small beads mill (QIAGEN K.K., Tokyo, Japan). These disrupted samples were preserved at −80°C until used. Total RNA was extracted from the samples according to the manufacture’s protocol for the TRI Reagent. Extracted RNA was treated with 2% RNase-free DNase I (New England BioLabs Japan Inc., Tokyo, Japan) at 37°C for 40 minutes and purified by phenol/chloroform extraction and ethanol precipitation. We measured the concentration of the total RNA using a spectrophotometer (DS-11+, Denovix Inc., Wilmington, USA). fs-cDNA was synthesized from 350 ng of the total RNA using SuperScript III Reverse Transcriptase (Thermo Fisher Scientific K.K., Tokyo, Japan). We diluted the fs-cDNA to 1:2 with MilliQ water and preserved it at −30°C until it was used in RT-qPCR assay. The RT-qPCR assays were performed using a LightCycler 96 instrument (Roche, Basel, Switzerland) according to the manufacture’s protocol with the THUNDERBIRD SYBR qPCR Mix (TOYOBO Co. Ltd., Osaka, Japan). The reaction volume was 10 µl. We used 1 µl of the fs-cDNA as templates. The preparation of the RT-qPCR solution proceeded on ice. The protocol of the RT-qPCR was as follows: preincubation at 95°C for 600 seconds and 45 cycles of three-step reactions, such as denaturation at 95°C for 15 seconds, annealing at 60°C for 15 seconds and extension at 72°C for 45 seconds. We used *ribosomal protein 49* (*rp49*) as a reference gene, as described by *Ohde et al. (2011)*. We designed primer sets of the target genes by the Primer3Web version 4.1.0 (*Untergasser et al., 2012*) following the manufacture’s recommended condition of the THUNDERBIRD SYBR qPCR Mix. We confirmed the primers’ specificity using melting curves ranging from 65°C to 95°C. We selected primer sets exhibiting a single peak. The primers are listed in Table 6. Each RT-qPCR was technically replicated three times. Some samples were excluded before analyzing the data when the Ct value of any genes was not detected (ND) in one or more replicates or when the Ct value of the reference gene deviated from that of other samples. In these removed data, a technical error was suspected. We calculated the expression level of target genes by the 2^-ΔΔCt^ method (*Livak and Schmittgen, 2001*) and performed the Brunner–Munzel (BM) test for ΔCt value. The BM test was carried out using R-v4.0.3. with the *brunnermuzel.test* function of the “brunnermuzel” package (https://cran.r-project.org/web/packages/brunnermunzel/index.html). Holm’s method was used for multiple comparison analyses between the control and treatments. The data are listed in Table 2. Also, its source data can be found in Table2–Source Data 1. In the *dsx* expression of the RNAi male, we performed the Smirnov-Grubbs (SG) test for ΔCt value using the *grubbs.test* function of the “outliers” package in R (https://cran.r-project.org/web/packages/outliers/index.html) (Table 3). An outlier was detected in the *dsx* RNAi male. We repeatedly performed the SG test using the data excluding the outlier. No further outliers were detected. Lastly, we re-analyzed the data, excluding the outlier, using the BM test (Table 2).

**Table 3.** Results of Smirnov–Grubbs’ test for expression level of *dsx* mRNA in nymphal RNAi males. The determination of whether a value is an outlier or not is based on the *P*-value and is shown in the “outlier?” column.

| data | max/min | value | *G* | *U* | *P*-value | outlier? |
| --- | --- | --- | --- | --- | --- | --- |
| *egfp* | max | 10.45 | 1.8587 | 0.65738 | 2.73.E-01 | no |
|  | min | 5.31 | 1.59672 | 0.74715 | 5.74.E-01 | no |
| *dsx* | max | 10.47 | 0.97 | 0.85 | 1.00.E+00 | no |
|  | min | 5.23 | 2.27 | 0.15688 | 5.14.E-03 | yes |
| *dsx-like* | max | 7.70 | 1.01 | 0.88 | 1.00.E+00 | no |
|  | min | 5.12 | 2.08 | 0.47 | 8.35.E-02 | no |
| reanalysis of<br>*dsx* | max | 10.47 | 1.52 | 0.55 | 3.52.E-01 | no |
|  | min | 8.44 | 1.42 | 0.61 | 4.70.E-01 | no |

### RNAi analysis

Nymphal RNAi can be used to analyze the roles of genes during postembryonic development. The sexual differentiation of insects is generally assumed to be a cell-autonomous mechanism that is independent of systemic hormonal-control (*Verhulst and van de Zande, 2015*) as discussed in *De Loof and Huybrechts (1998)*and *Bear and Monteiro (2013)* and progresses during postembryonic development. Therefore, nymphal RNAi is the most effective tool to investigate the roles of genes on sexual trait formation. To reduce the possibility of off-target effects, the dsRNA was designed to avoid the region of the DM domain. We also confirmed that the dsRNA had no contiguous matches of more than 20 bases with other regions of the genome. To produce templates for the dsRNA, we cloned the regions of *dsx* and *dsx-like* from the fs-cDNA using the same method as the RACE analysis. We amplified the template DNAs from purified plasmids with PCR using Q5 High-Fidelity DNA Polymerase and purified the amplified DNA with the phenol/chloroform extraction and the ethanol precipitation. dsRNA was synthesized from the purified DNA using Ampliscribe T7-Flash Transcription kits (Epicentre Technologies, Co., Wisconsin, USA). We designed the PCR primers using the Primer3Web version 4.1.0 (*Untergasser et al., 2012*). The PCR primers are listed in Table 6. In nymphal RNAi analysis, we injected the dsRNAs repeatedly into the abdomen of the nymphs of *T. domestica* with each molt from the fourth or fifth instar to thirteenth instar to sustain the RNAi effect during postembryonic development. The initial stage was the same within a single experiment. This repeated nymphal RNAi was effective in some insects such as *Blattella germanica* (*Wexler et al., 2014*). We sampled the individuals one, three, and five days after molting, using phenotypic observations, analysis of *dsx* knockdown effects, and the oocyte size and number. To determine the sex of individuals, we initially observed the gonads: testis and ovary. In our nymphal RNAi analysis, the gonads completely formed and there was no difference between the control and *dsx* RNAi individuals in external morphology (Figures 1–figure supplement 3, Figure2–figure supplement 1A). Therefore, at least in this assay, the gonad morphology was effective for the identification of the sex of *dsx* RNAi individuals. Individuals with testis were males and those with ovaries were females. For the analysis of *vtg* genes, we used the adult RNAi assay. *T. domestica* molts throughout its life, even after sexual maturation, and produces *vtg* during each adult instar (*Rouset and Bitsch, 1993*). In the adult RNAi analysis, we injected dsRNA repeatedly into the adults every three days. The dsRNA was initially injected into adults 12 hours after molting. We sampled the adults at 720±20 minutes after subsequently molts, to analyze the *vtg* mRNA levels.

### Phenotype observation

We dissected thirteenth instar individuals in PBS using tweezers and removed the thoraxes, reproductive systems, and external genital organs. We took images using the digital microscope system (VHX-5000, KEYENCE, Tokyo, Japan). The thoraxes and external genital organs were fixed with FAA fixative (formaldehyde: ethanol: acetic acid = 15:5:1) at 25°C overnight and then preserved in 90% ethanol. We used the length of the prothorax as an indicator of body size. To measure the prothoracic width, the prothoracic notum was removed from the fixed thorax after treatment with 10% NaOH solution at 60°C for 30 minutes to dissolve the soft tissues. The notum was mounted in Lemosol on a microscope slide. The prepared specimens were imaged using a KEYENCE VHX-5000. With the microscope at 50×, the length of the notum was measured. The ovipositor length and oocyte size were also measured using the microscope at 20× and 50×. The oocyte size was taken to be the major length of the late vitellogenic oocyte at the posterior most part of the ovariole. To count the sperm number, sperm was collected from seminal vesicles and diluted with 5 ml MilliQ water. 50 µl of the diluted sperm was spotted on a microscope slide and dried overnight. We technically replicated the measurement three times for ovipositor length and six times in sperm number and calculated these means. Measurement was performed by blinding the treatment. We counted the number of oocytes in ovarioles using an optical microscope at 50× (Olympus, Tokyo, Japan). A generalized linear model (GLM) was used to analyze differences in ovipositor length and oocyte size (length data) and sperm and oocyte number (count data) among RNAi treatments. The body size, target genes, and interactions between the target genes were used as explanatory variables. The length was assumed to follow a Gaussian distribution, and the count data to have a negative binomial distribution. We used R-v4.0.3 in these analyses and the *glm* and the *glm.nb* (MASS package) functions for the length and count data, respectively. To analyze the contribution of the explanatory variables, a likelihood ratio test for the result of GLM was performed using the *Anova* function of the car package. The statistical results are listed in Tables 4 (female) and 5 (male). Also, the source data are reported in Table 4–source data 1 and Table 5–source data 1.

**Table 4.** Results of generalized linear model of female traits. Significant levels are indicated by asterisks in significance column: \**P*<0.05, \*\**P* < 0.01, \*\*\**P* < 0.001. n.s. means non-significance.

| objective variable | sample size (N) | explanatory variables | LR Chisq | Df | P-value | significance | Fig # |
| --- | --- | --- | --- | --- | --- | --- | --- |
| ovipositor length | *egfp*: 10, *dsx* all: 17,<br>*dsx-like*: 9, *dsx+dsx-like*: 12, total: 48 | *dsx* RNAi | 0.26 | 1 | 6.08.E-01 | n.s. | Figure 2H |
|  |  | *dsx-like* RNAi | 0.58 | 1 | 4.47.E-01 | n.s. |  |
|  |  | prothoracic width | 297.00 | 1 | 2.00.E-16 | *** |  |
|  |  | *dsx* RNAi:*dsx-like* RNAi | 2.67 | 1 | 1.02.E-01 | n.s. |  |
| ovipositor length | *egfp*: 8, *dsx* all: 8,<br>*dsx male-type*: 7,<br>*dsx female-type*: 13,<br>total: 36 | *dsx* all RNAi | 1.29 | 1 | 2.56.E-01 | n.s. | Figure 2–figure<br>supplement 3 |
|  |  | *dsx male-type* RNAi | 1.29 | 1 | 2.56.E-01 | n.s. |  |
|  |  | *dsx female-type* RNAi | 0.42 | 1 | 5.18.E-01 | n.s. |  |
|  |  | prothoracic width | 146.52 | 1 | 2.00.E-16 | *** |  |
| prothoracic width | *egfp*: 10, *dsx* all: 17,<br>*dsx-like*: 9, *dsx+dsx-like*: 13, total: 49 | *dsx* RNAi | 2.17 | 1 | 1.41.E-01 | n.s. | Figure 2–figure<br>supplement 4 |
|  |  | *dsx-like* RNAi | 1.96 | 1 | 1.61.E-01 | n.s. |  |
|  |  | *dsx* RNAi:*dsx-like* RNAi | 2.16 | 1 | 1.41.E-01 | n.s. |  |
| previtellogenic<br>oocyte number | *egfp*: 7, *dsx* all: 16,<br>*dsx-like*: 10,<br>*dsx+dsx-like*: 9, total:<br>42 | *dsx* RNAi | 0.08 | 1 | 7.83.E-01 | n.s. | Figure 3D |
|  |  | *dsx-like* RNAi | 0.23 | 1 | 6.31.E-01 | n.s. |  |
|  |  | prothoracic width | 0.97 | 1 | 3.25.E-01 | n.s. |  |
|  |  | *dsx* RNAi:*dsx-like* RNAi | 0.06 | 1 | 8.10.E-01 | n.s. |  |
| early vitellogenic<br>oocyte number | *egfp*: 7, *dsx* all: 16,<br>*dsx-like*: 10,<br>*dsx+dsx-like*: 9, total:<br>42 | *dsx* RNAi | 0.11 | 1 | 7.45.E-01 | n.s. | Figure 3D |
|  |  | *dsx-like* RNAi | 0.08 | 1 | 7.77.E-01 | n.s. |  |
|  |  | prothoracic width | 3.50 | 1 | 6.14.E-02 | n.s. |  |
|  |  | *dsx* RNAi:*dsx-like* RNAi | 0.62 | 1 | 4.30.E-01 | n.s. |  |
| late vitellogenic<br>oocyte number | *egfp*: 7, *dsx* all: 16,<br>*dsx-like*: 10,<br>*dsx+dsx-like*: 9, total:<br>42 | *dsx* RNAi | 0.46 | 1 | 5.00.E-01 | n.s. | Figure 3D |
|  |  | *dsx-like* RNAi | 1.12 | 1 | 2.90.E-01 | n.s. |  |
|  |  | prothoracic width | 9.84 | 1 | 1.71.E-03 | ** |  |
|  |  | *dsx* RNAi:*dsx-like* RNAi | 2.16 | 1 | 1.42.E-01 | n.s. |  |
| oocyte size | *egfp*: 5, *dsx* all: 9,<br>*dsx-like*: 7, *dsx+dsx-*<br>*like*: 2, total: 23 | *dsx* RNAi | 0.1689 | 1 | 6.81.E-01 | n.s. | Figure3–figure<br>supplement 3 |
|  |  | *dsx-like* RNAi | 3.4291 | 1 | 6.41.E-02 | n.s. |  |
|  |  | prothoracic width | 4.273 | 1 | 3.87.E-02 | * |  |
|  |  | *dsx* RNAi:*dsx-like* RNAi | 1.4509 | 1 | 2.23.E-01 | n.s. |  |

The source data of this table can be obtained in the Table4–source data 1 file.

**Table 5.**
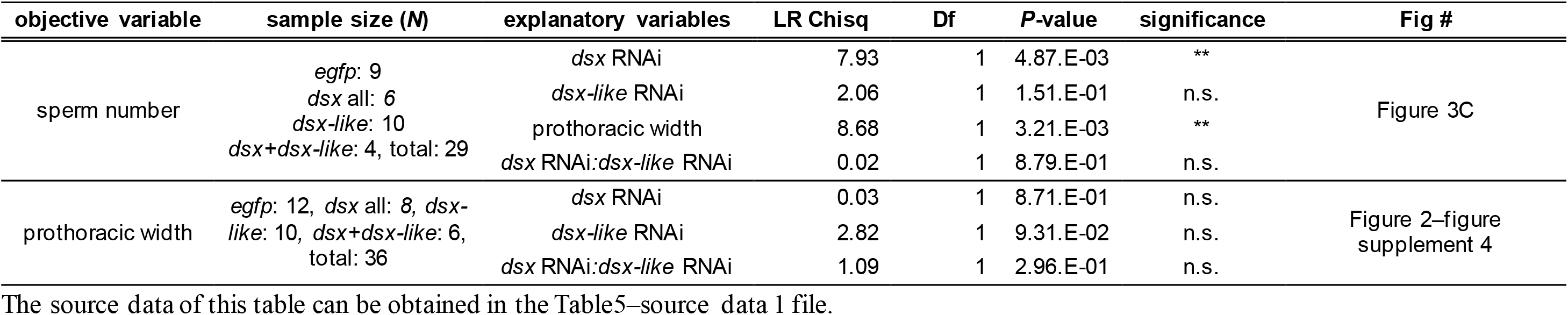
Results of generalized linear model of male traits. *P*-values are indicated by asterisks in significance column: \*\**P* < 0.01. n.s. means non-significance.

**Table 6.**
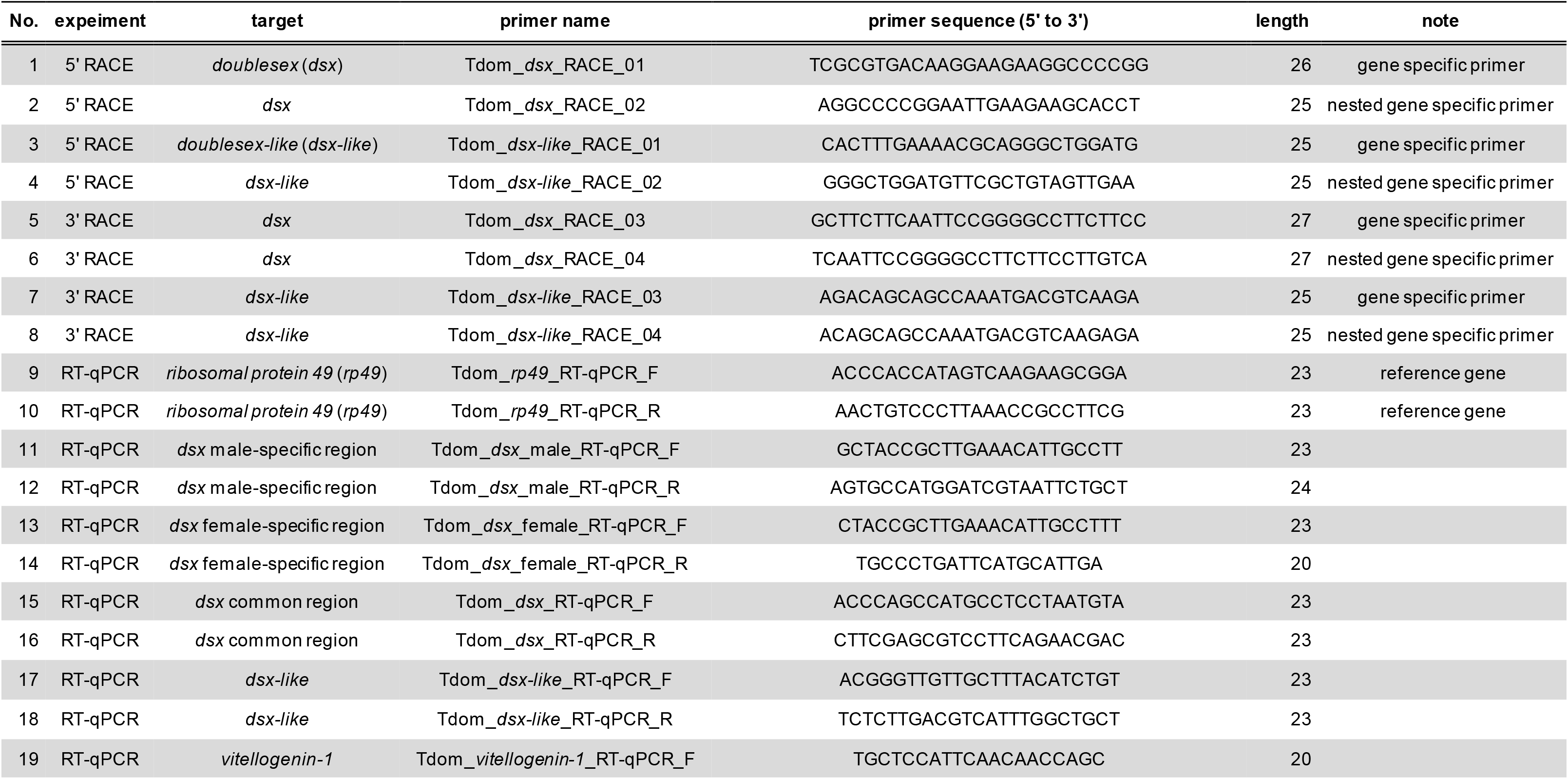
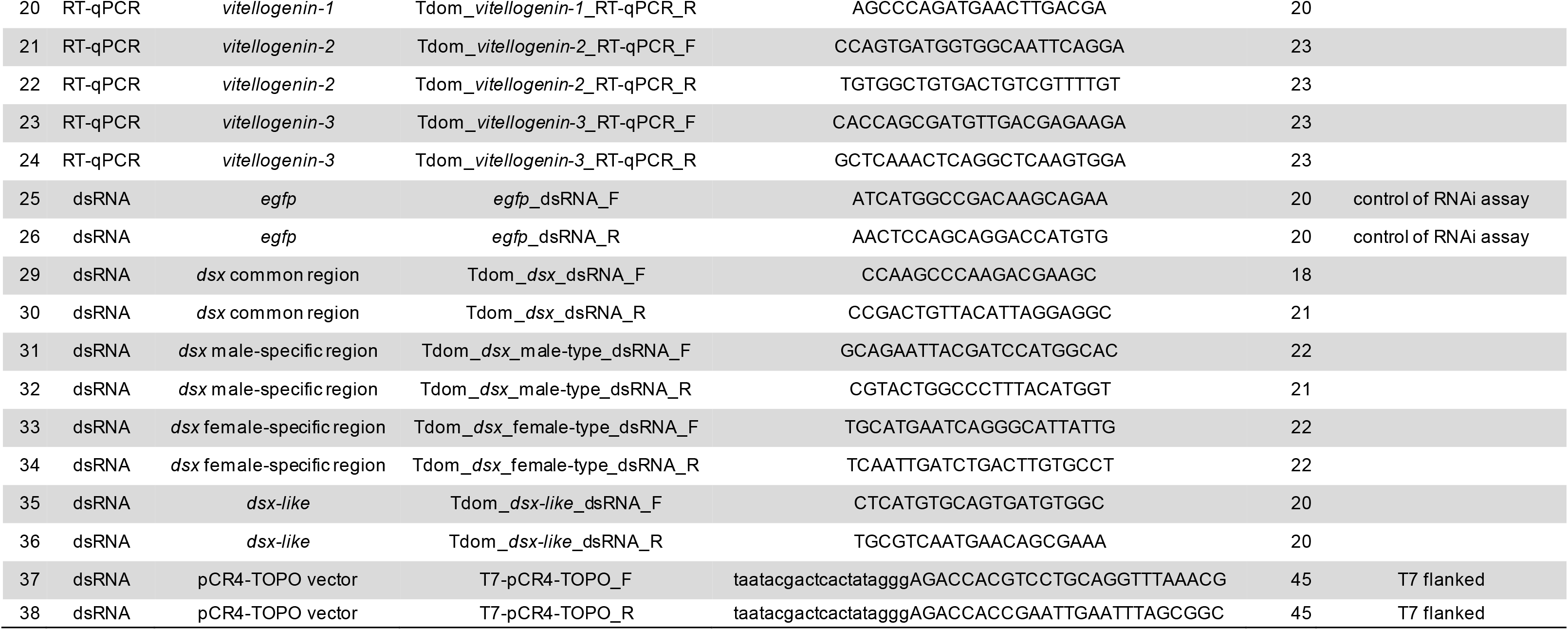
Primers’ list used in this study.

### Scanning Electron Microscopy (SEM)

The NanoSuit method (*Takaku et al., 2013*) was used for the SEM analysis. Male penises and female ovipositors preserved in 90% ethanol were washed with distilled water and immersed in 1% Tween20 at 25°C for 10 minutes. The samples were mounted on stubs and imaged using a low-vacuum SEM (DX-500; KEYENCE, Tokyo, Japan).

### Histology

The gonads of RNAi individuals were fixed with Bouin’s fixative (saturated picric acid: formaldehyde: glacial acetic acid = 15:5:1) at 25°C overnight and washed with 90% ethanol plus Lithium Carbonate (Li_2_CO_3_). The ovipositors of RNAi individuals were fixed with FAA fixative at 25°C overnight and then were transferred into 90% ethanol. The samples were dehydrated and cleared with an ethanol-butanol series. The cleared samples were immersed and embedded in paraffin at 60°C. The paraffin blocks were polymerized at 4°C and cut into 5 µm thick sections using a microtome (RM2155: Leica, Wetzlar, Germany). The sections were mounted on microscope slides coated with egg white-glycerin and stained using Delafield’s Hematoxylin and Eosin staining. The stained slides were enclosed with Canada balsam. We observed the slides on an optical microscope (Olympus, Tokyo, Japan) and took photos using a digital single-lens reflex camera (Nikon, Tokyo, Japan).

### Comparison of exon-intron structure and amino acid sequences

We obtained exon-intron structures of *dsx* in seven insect species from NCBI. The female- and male-specific regions were visualized as red and blue colors, respectively. The exon-intron structures were mapped on a phylogenetic hypothesis based on *Misof et al. (2014)*. We obtained amino acid sequences of Dsx isoforms of 10 species from the NCBI database. *Wexler et al. (2014)* showed that *dsx* of *Pediculus humanus* (Psocodea) has isoforms without sex-specificity. In this study, based on the blast search and exon structure, we determined that PhDsx1 in *Wexler et al. (2019)* corresponded to the *dsx* female-type and PhDsx2 to *dsx* male-type. The sequences were aligned using MAFFT version 7 with a -linsi option.

### Data availability

The draft genome data was deposited in the DNA Data Bank of Japan (Accession number: DRA005797; Bioproject: PRJDB5781). The raw read data of the transcriptome was in the NCBI Sequence Read Archive (Accession numbers: SRR13870115–SRR13870124; Bioproject: PRJNA707122). The sequences of *dsx* male-type, *dsx* female-type, and *dsx-like* are also in GenBank (Accession numbers: MW711323, MW711324, and MW711325, respectively).

## Acknowledgments

We would like to thank Dr. Takahiro Ohde (National Institute for Basic Biology; Kyoto University) for his help to extract genomic DNA of *T. domestica* and for a great advice to the discussion. We are also grateful to Dr. Toshiya Ando, Dr. Taro Nakamura, Dr. Shinichi Morita, Dr. Hiroki Sakai, and Dr. Tatsuro Konagaya (National Institute for Basic Biology) for technical advice and discussion on this manuscript. We also express our gratitude to Dr. Morita for providing a photo of *Daphnia magna*. Computations were performed on the NIG supercomputer at ROIS, National Institute of Genetics and the Data Integration and Analysis Facility, National Institute for Basic Biology. We thank the Model Plant Research Facility, NIBB Bioresource Center for providing the network camera system. This work was supported by the JSPS KAKENHI Grant numbers JP25660265, JP16H02596, and JP16H06279 (PAGS) for TN and the Sasakawa Scientific Research Grant from The Japan Science Society for YC.

## Author Contributions

YC and TN conceived this study. YC performed the cloning of *dsx* and *dsx-like*, the RNAi, the RNA-seq, and the molecular phylogenetic analysis. AT sequenced the genome. MO and TI performed the de novo genome assembly. YC and T. wrote this manuscript; all authors commented on the manuscript.

## Video

**Video 1. Time-lapse imaging of molting of *Thermobia domestica*.**

## Figure Supplements

**Figure 1–figure supplement 1.**
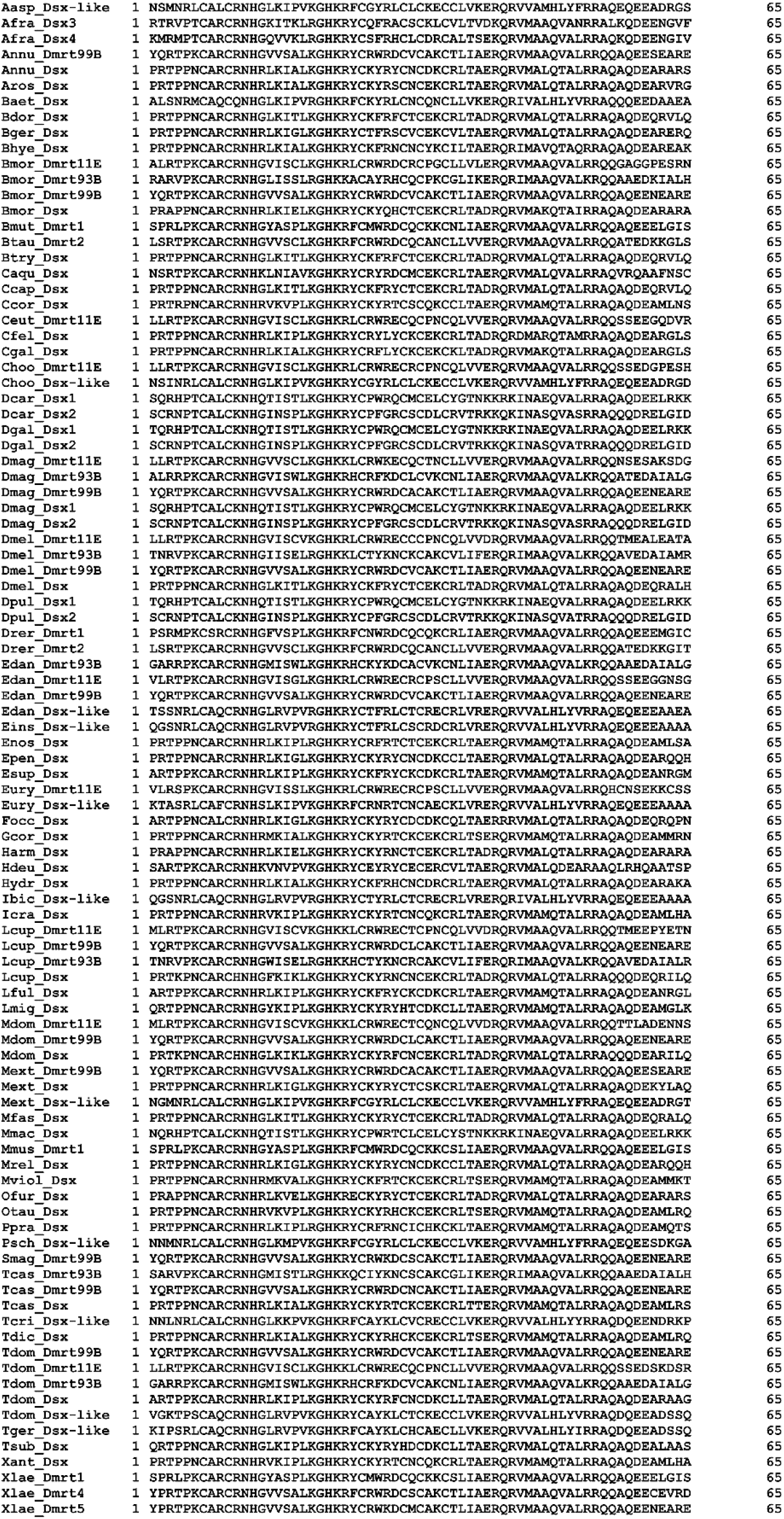
Multiple sequence alignment of DM domain of DMRT family proteins for molecular phylogenetic analysis. The 65 amino acids of 97 DMRT proteins were used for the molecular phylogenetic analysis.

**Figure 1–figure supplement 2.**
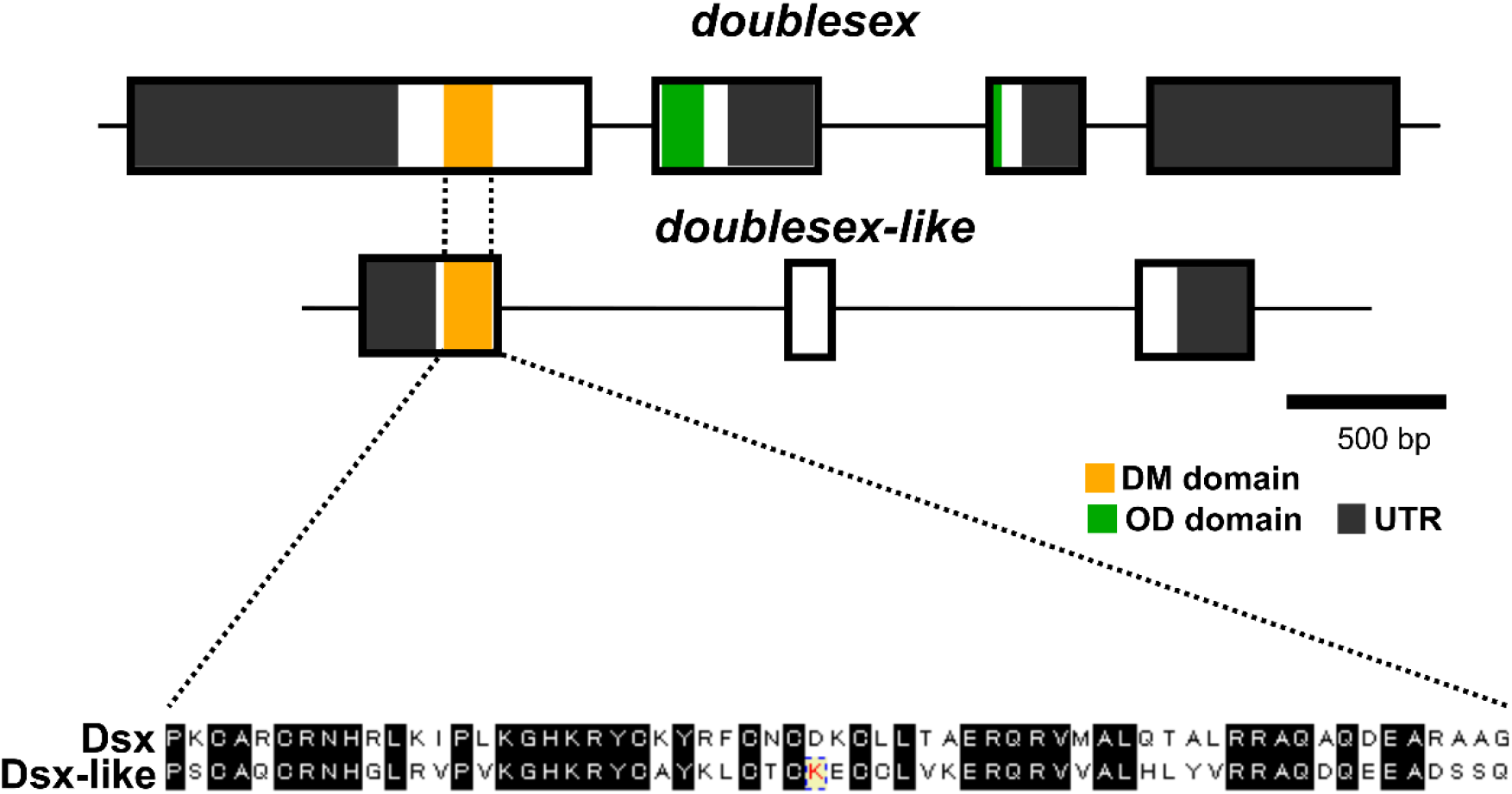
Comparison of DM domain sequences between Dsx and Dsx-like. The upper figures show the gene structures of *dsx* and *dsx-like*, and the lower one is the result of the multiple sequence alignment of the DM domain in Dsx and Dsx-like of *Thermobia domestica*. The black-highlighted sequences are shared in both proteins.

**Figure 1–figure supplement 3.**
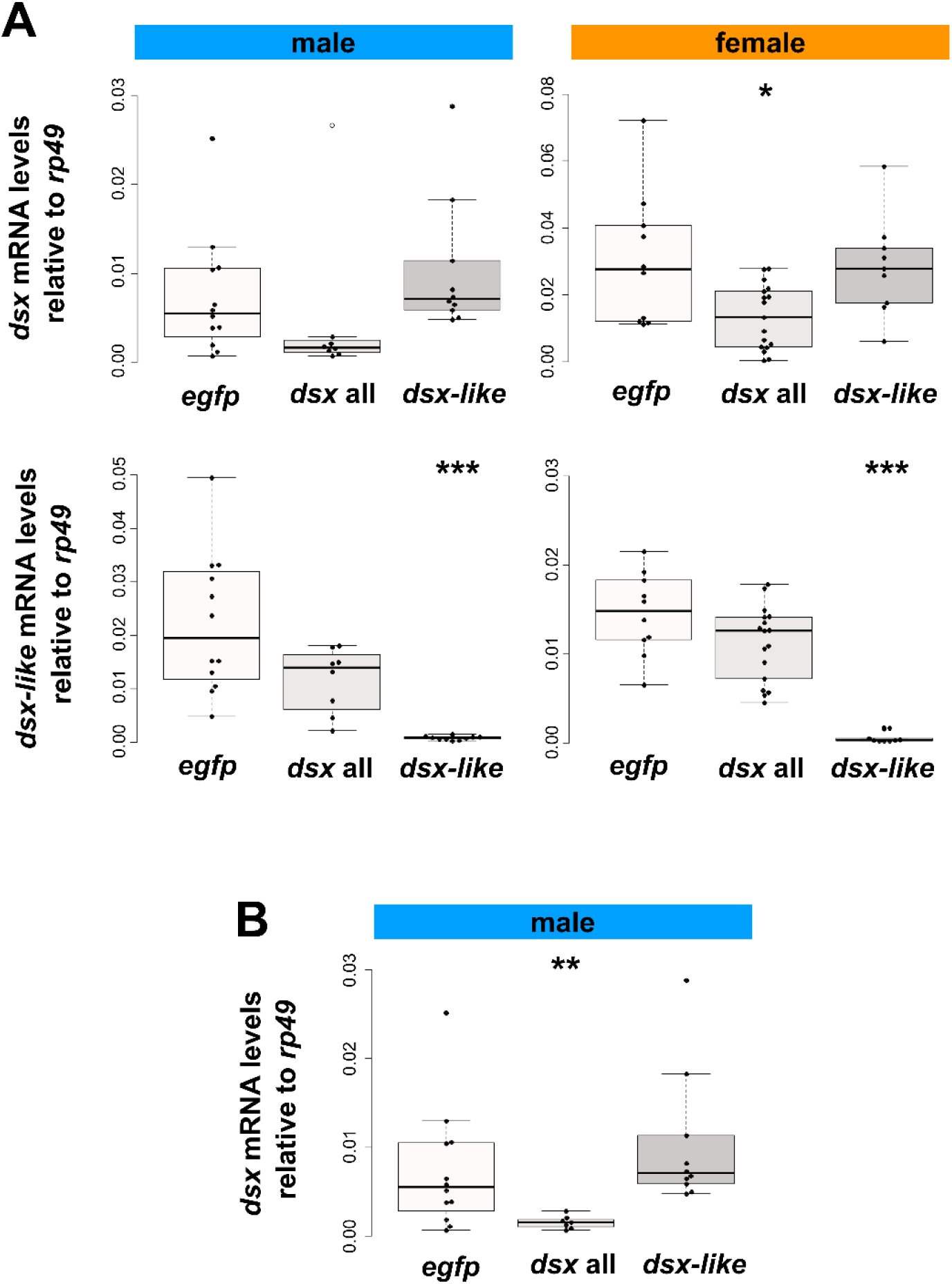
Expression of *dsx* and *dsx-like* mRNA in nymphal RNAi individuals. (A) results of RT-qPCR assay. The expression profiles of *dsx* and *dsx-like* mRNA were analyzed by RT-qPCR assay and are indicated by their expression level relative to the expression of a reference gene (*ribosomal protein 49*). The upper graphs are the expression of *dsx* mRNA and the lower ones are that of *dsx-like* mRNA. The left column is the result in males and the right one is that in females. (B) the expression level of *dsx* mRNA in the nymphal RNAi males after excluding an outlier. To test the outlier, the Smirnov–Grubbs’ test was performed. Results of the Smirnov–Grubbs’ test are shown in Table 3. Results of the Brunner–Munzel test are indicated by asterisks: \**P* < 0.05; \*\**P* < 0.01; \*\*\**P* < 0.001 and is also described in Table 2. *P* > 0.05 is not shown. Each plot is an individual. White plot is the outlier. Sample size are listed in Table 2.

**Figure 2–figure supplement 1.**
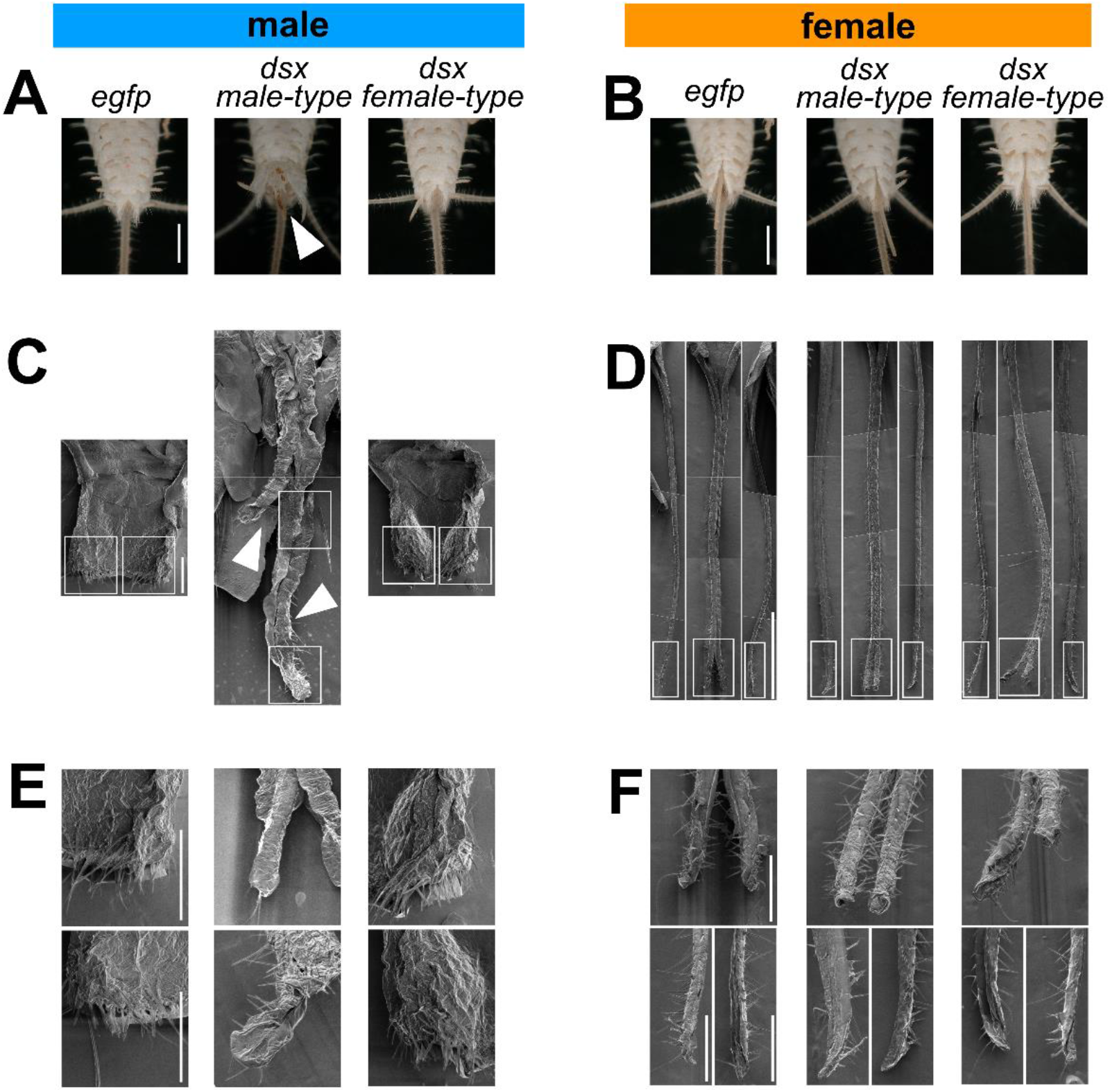
External genital organs of nymphal RNAi individuals. (A) The penis in ventral view of *dsx* male or female-type RNAi males. (B) The ovipositor in ventral view of *dsx* male or female-type RNAi females. (C) SEM images of the male penis or the ovipositor-like organ. White frames are the areas shown in (E). (D) SEM images of the female ovipositor. Each image is separated into three parts. The left and right image is each lobe of the valvula I. The middle image is the valvula II. White frames are the areas showed in (F). (E) Higher magnification SEM images of the male genital organ in (C). Each image shows the higher magnification view of the area enclosed by the white frame in (C). (F) Higher magnification SEM images of the ovipositor. Each image shows the higher magnification view of the area enclosed by the white frame in (*D*). The arrowheads show the ovipositor-like organ in the *dsx* male-type RNAi male. Scales: 1000 µm (A, B, and D), 100 µm (C, E, and F).

**Figure 2–figure supplement 2.**
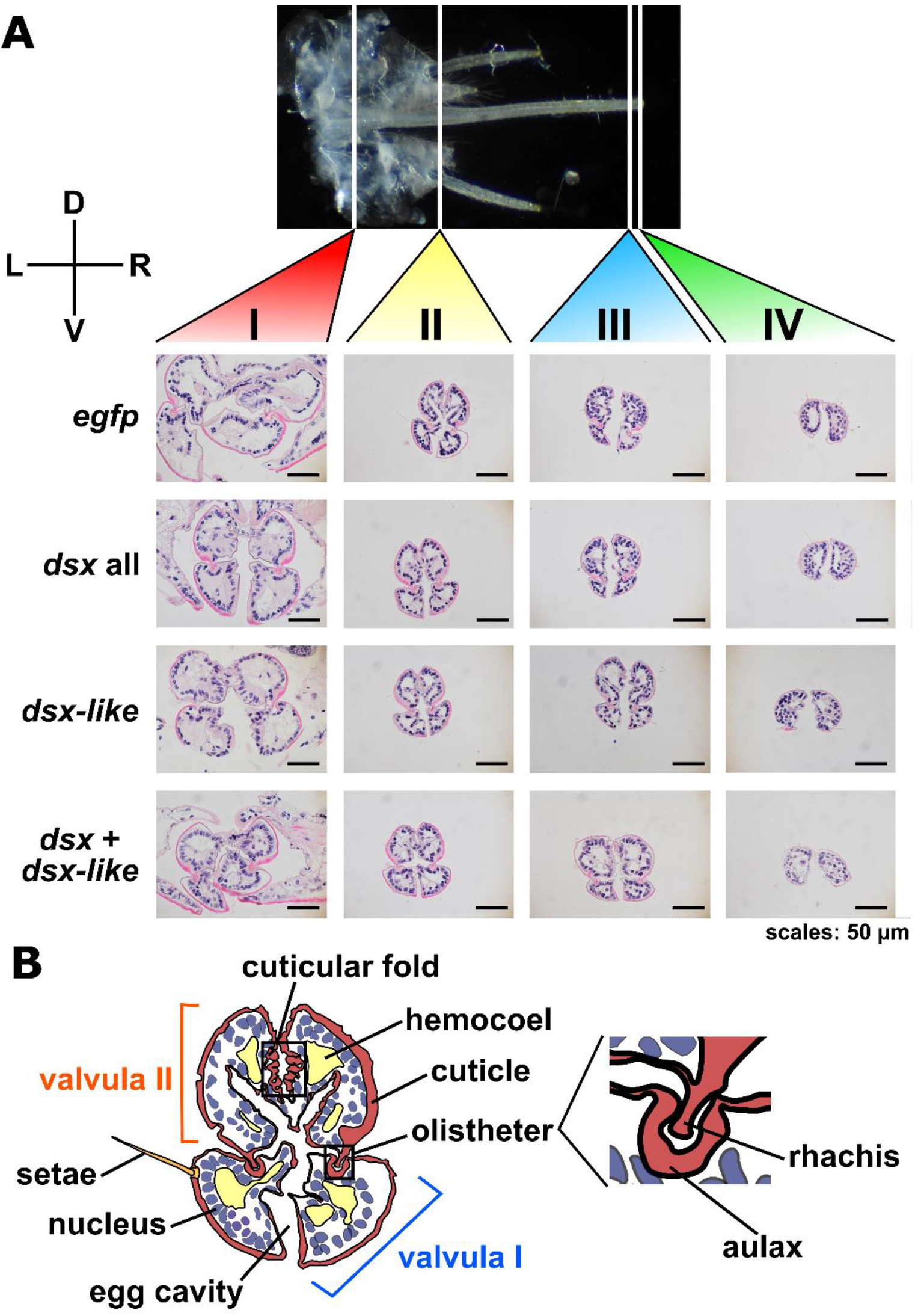
Morphology of ovipositor in nymphal RNAi individuals. (A) Cross-section of the ovipositor. The photos show the morphology of the ovipositor in four parts: I (proximal part), II (middle part), III (distal part), and IV (most-distal part). D, dorsal; L, left; R, right; V, ventral. Scales: 50 µm. (B) Schematic figure of the ovipositor morphology. This figure is based on the cross-section of the part II in the control female.

**Figure 2–figure supplement 3.**
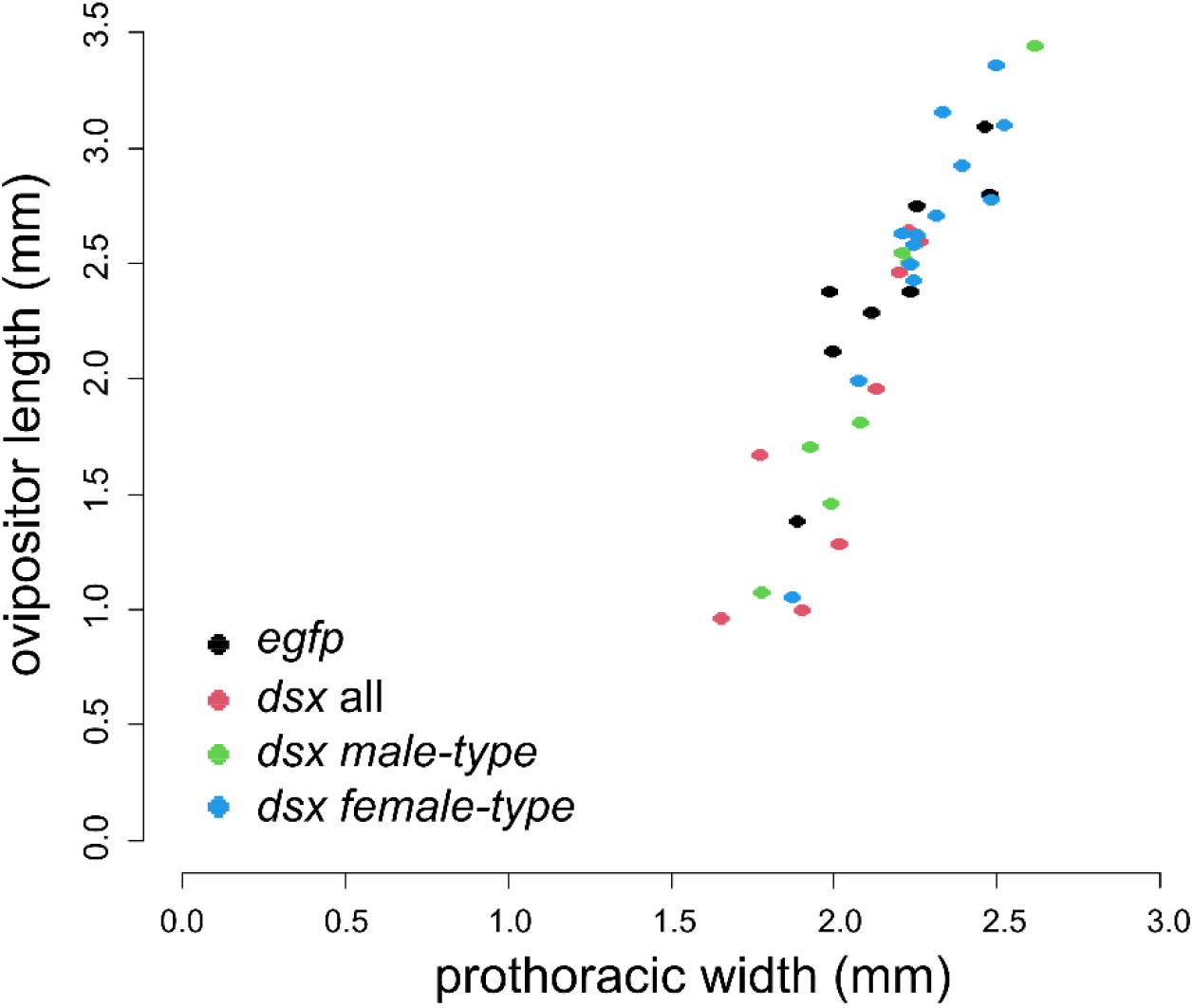
Ovipositor length of *dsx* isoforms RNAi individuals. The ovipositor length of *dsx* all, male-, and female-type RNAi females is plotted against the prothoracic width. The results of the statistical analysis are described in Table 4.

**Figure 2–figure supplement 4.**
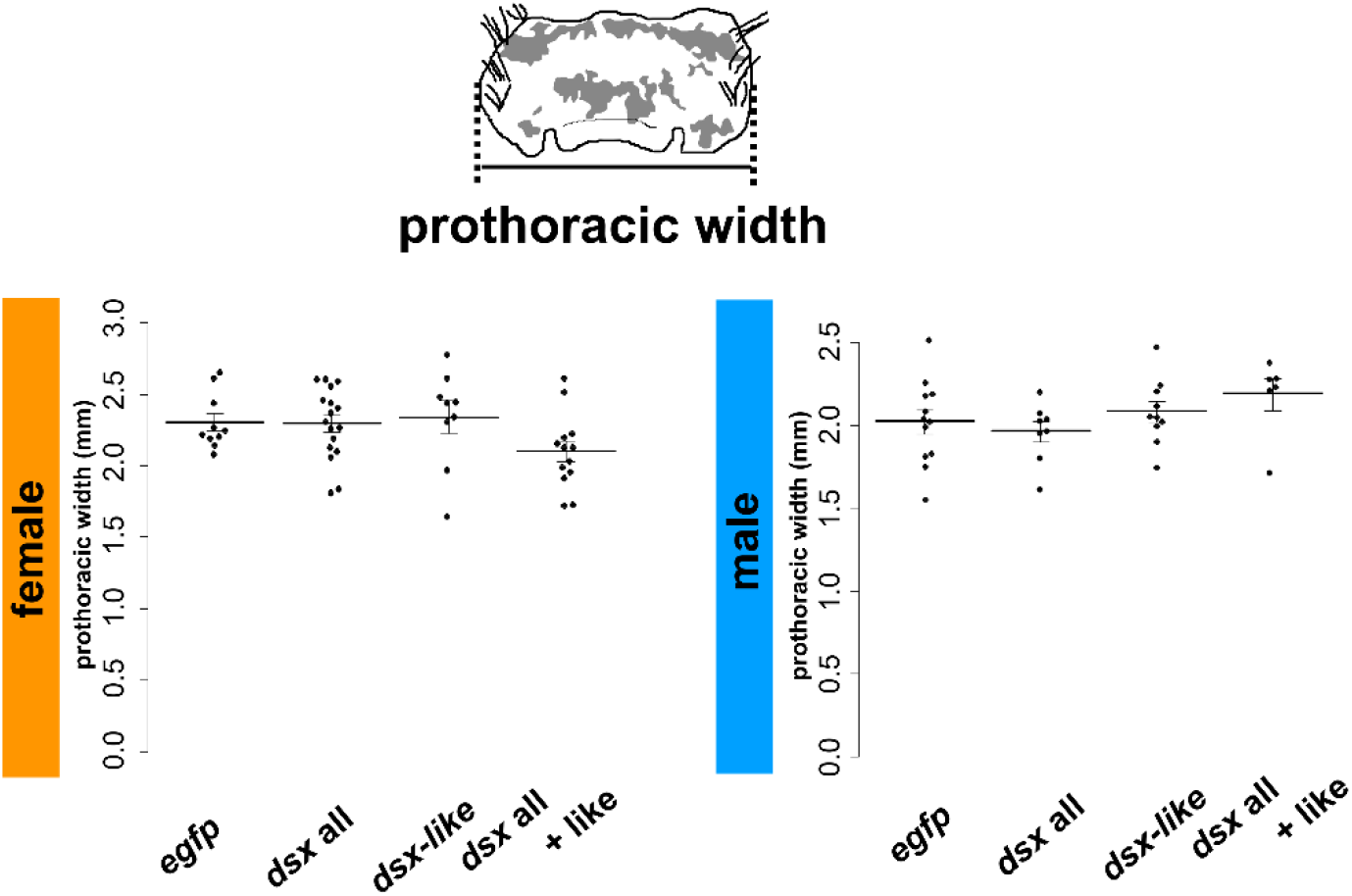
Prothoracic width of nymphal RNAi individuals. The left and right graphs show the female and the male size, respectively. Data shows mean±Standard Error (SE). The results of the statistical analysis are described in Tables 4 (female) and 5 (male).

**Figure 3–figure supplement 1.**
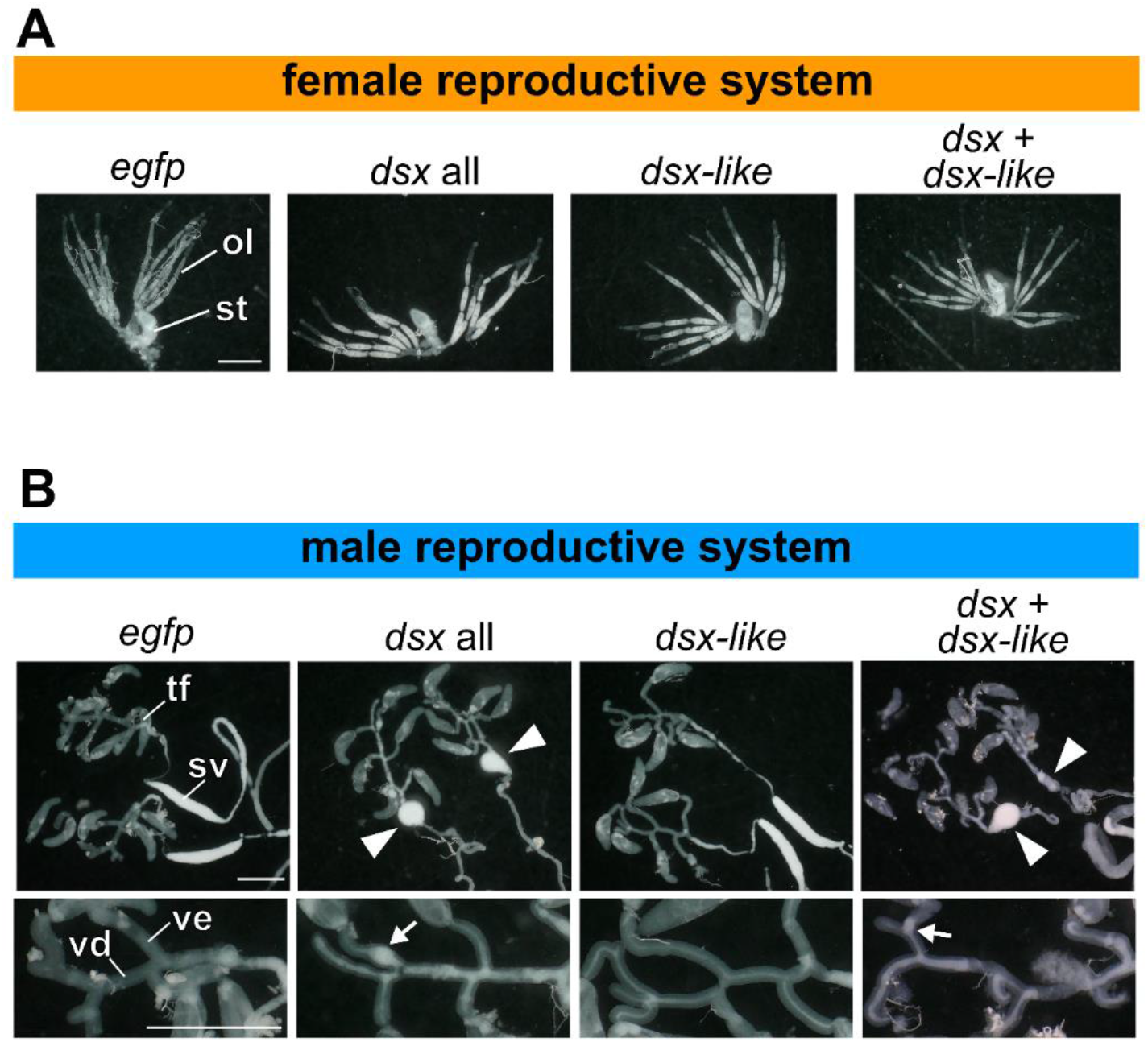
Gonads of nymphal RNAi individuals. (A) female reproductive system. ol, ovariole; st, spermatheca. (B) male reproductive system. The upper photos are the are from the testicular follicle to the seminal vesicle. The lower ones are the high-magnification image of the vas efferens and the vas deferens. The arrowheads show the round-shape seminal vesicle. The arrows indicate sperm clots in the vas efferens. sv, seminal vesicle; tf, testicular follicle; ve, vas efferens; vd, vas deferens. Scales: 1 mm.

**Figure 3–figure supplement 2.**
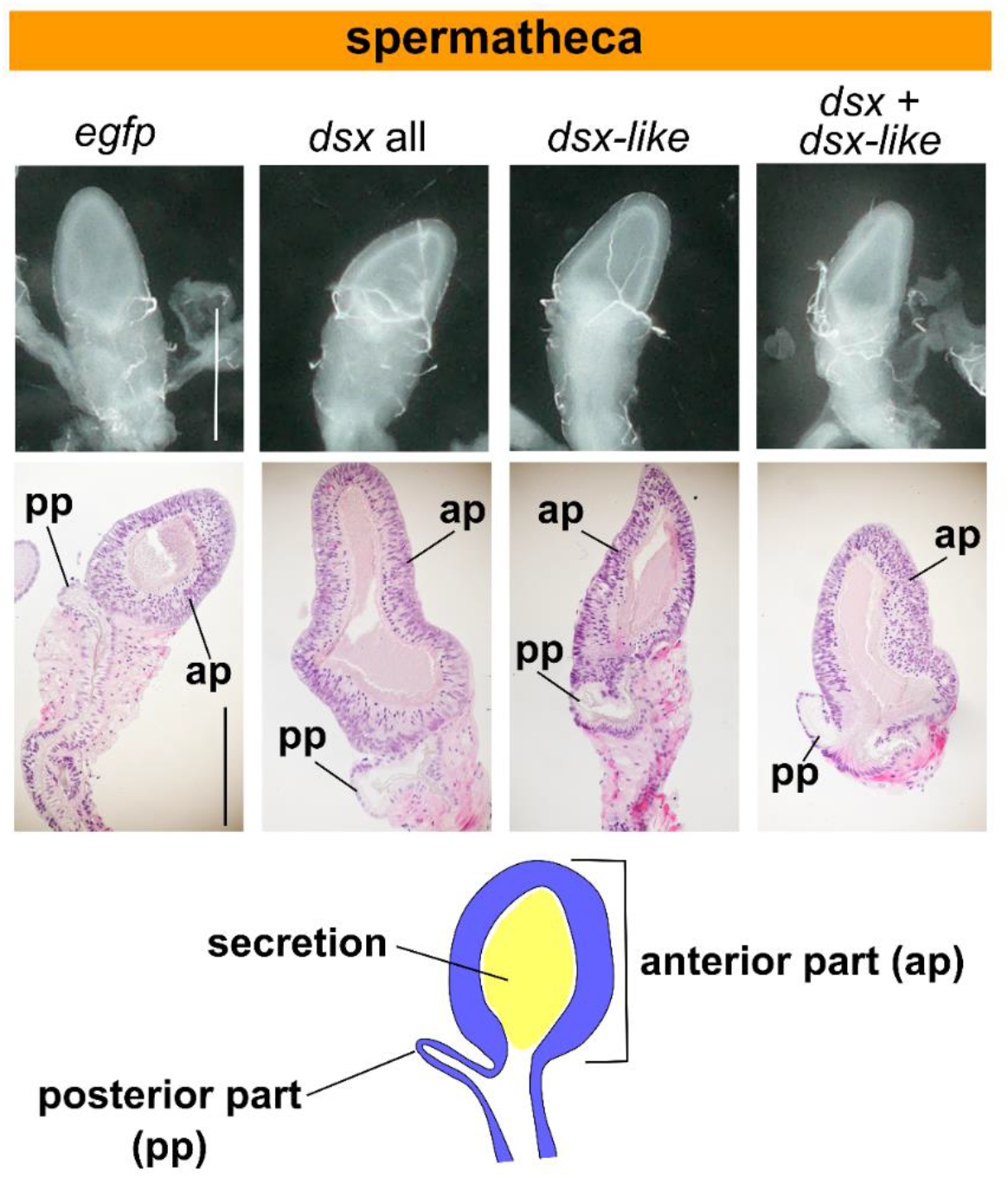
Morphology of spermatheca in nymphal RNAi individuals. The upper photos show the light microscopic images of the spermatheca. The middle ones are paraffin sections of the spermatheca. The lower one is the schematic image of the spermatheca of *T. domestica*. Scales: 500 µm.

**Figure 3–figure supplement 3.**
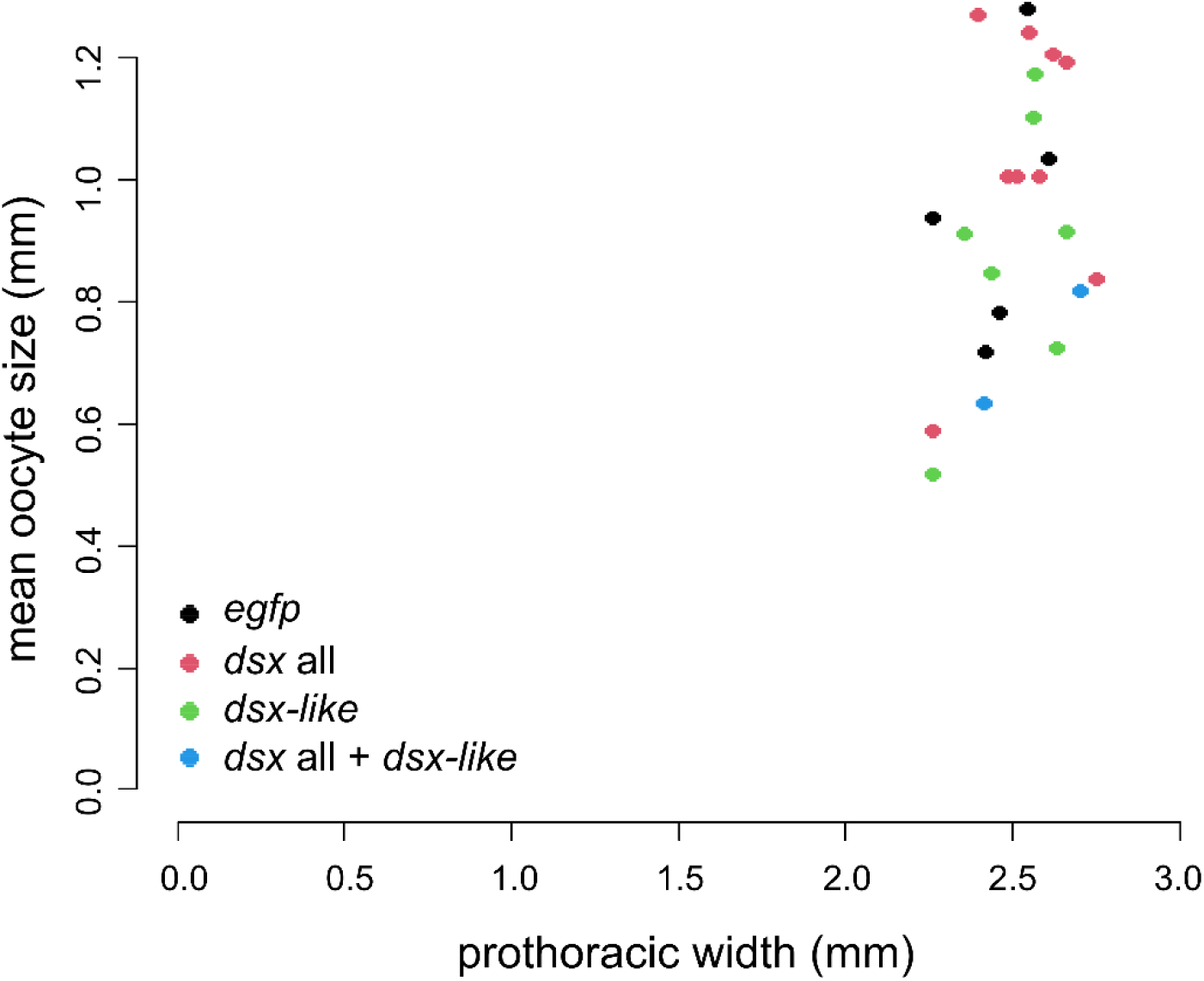
Oocyte size of *dsx* isoforms RNAi individuals. The oocyte size of *dsx* all, *dsx-like*, and both *dsx* and *dsx-like* RNAi females is plotted against the prothoracic width. The results of the statistical analysis are described in Table 4.

**Figure 4–figure supplement 1.**
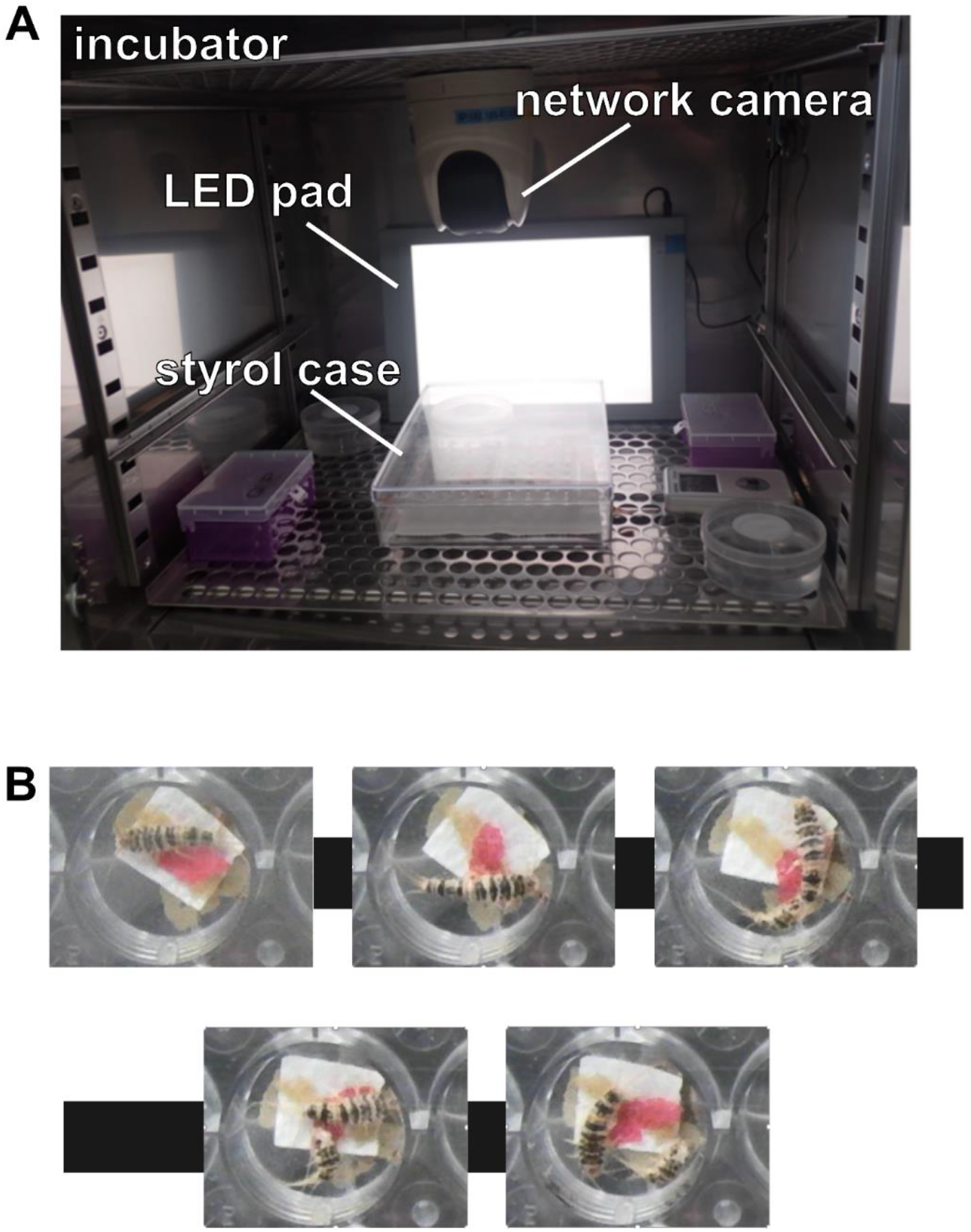
Time-lapse imaging system. (A) A photo of the time-lapse imaging system used to observe the molt of *T. domestica*. (B) The images during the molt.

**Figure 5–figure supplement 1.**
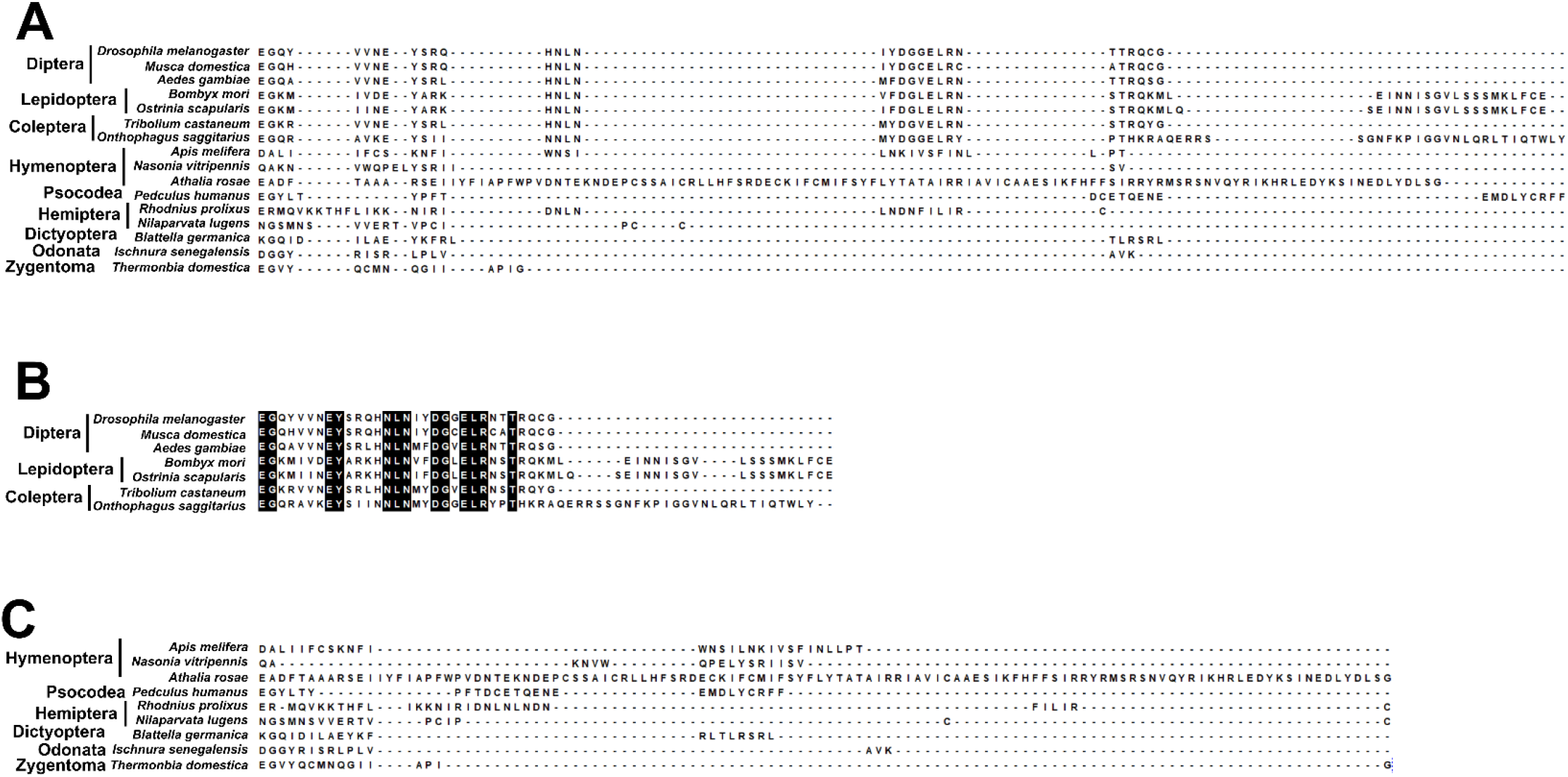
Multiple sequence alignments of insect Dsx female-specific region. (A) Comparison of the female-specific region of insect Dsx among the all taxa. (B) Comparison of the female-specific region of insect Dsx among the taxa with the dual-functionality. (C) Comparison of the female-specific region of insect Dsx among the taxa with the single-functionality. The black-highlighted sequences are shared in all given taxa.

**Figure 5–figure supplement 2.**
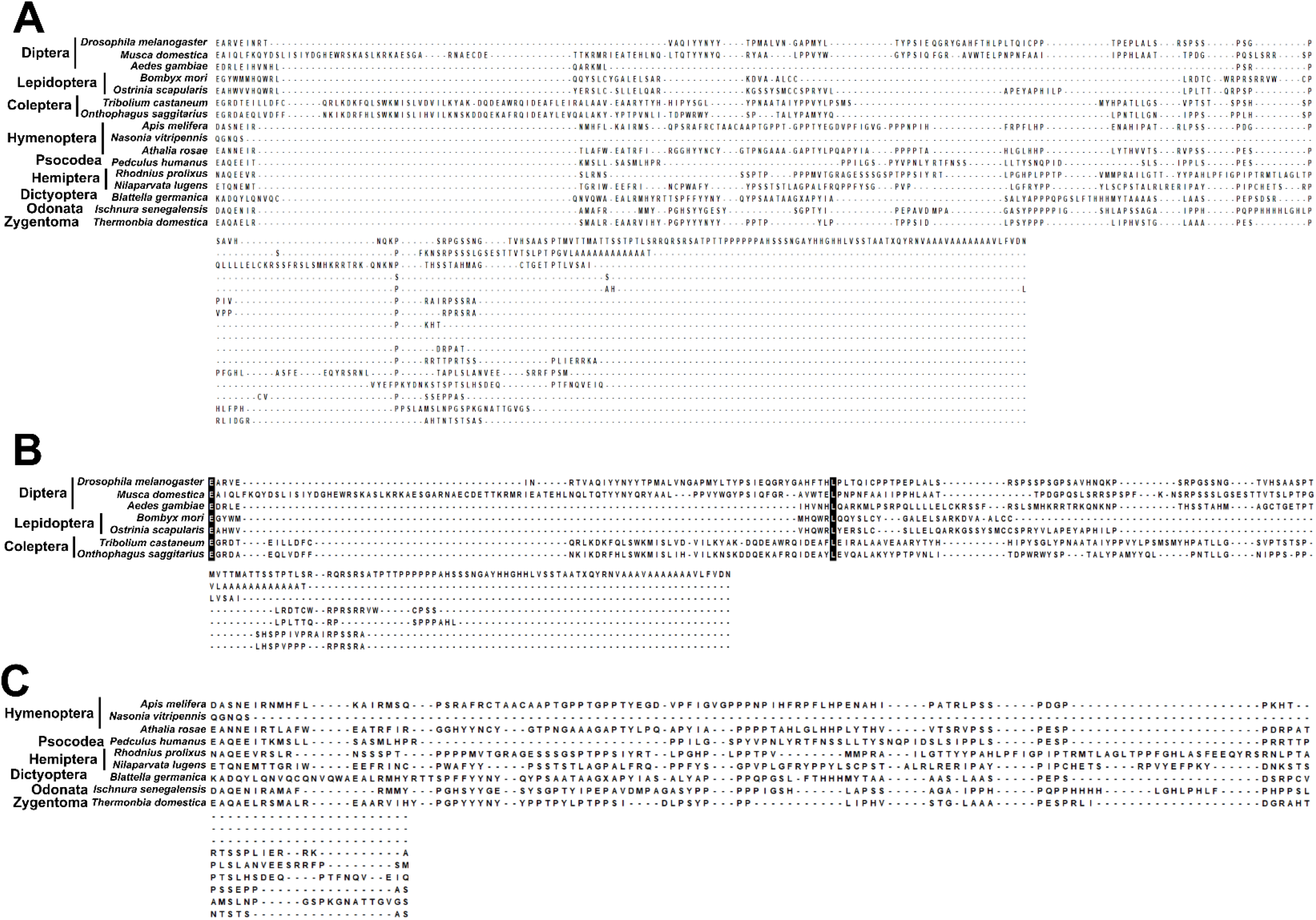
Multiple sequence alignments of insect Dsx male-specific region. (A) Comparison of the male-specific region of insect Dsx among the all taxa. (B) Comparison of the male-specific region of insect Dsx among the taxa with the dual-functionality. (C) Comparison of the male-specific region of insect Dsx among the taxa with the single-functionality. The black-highlighted sequences are shared in all given taxa.

